# RPA complexes in *Caenorhabditis elegans* meiosis; unique roles in replication, meiotic recombination and apoptosis

**DOI:** 10.1101/2020.06.27.174912

**Authors:** Adam Hefel, Nicholas Cronin, Kailey Harrel, Pooja Patel, Maria Spies, Sarit Smolikove

## Abstract

Replication Protein A (RPA) is critical complex that acts in replication and promotes homologous recombination by allowing recombinase recruitment to processed DSB ends. Most organisms possess three RPA subunits (RPA1, RPA2, RPA3) that form a trimeric complex critical for viability. The *Caenorhabditis elegans* genome encodes for RPA-1, RPA-2 and an RPA-2 paralog RPA-4. In our analysis, we determine that RPA-2 is critical for germline replication, and normal repair of meiotic DSBs. Interestingly, RPA-1 but not RPA-2 is essential for replication, contradictory to what is seen in other organisms, that require both subunits. In the germline, both RPA-1/2 and RPA-1/4 complexes form, but RPA-1/4 is less abundant and its formation is repressed by RPA-2. While RPA-4 does not participate in replication or recombination, we find that RPA-4 inhibit RAD-51 filament formation and promotes apoptosis on a subset of damaged nuclei. Altogether these findings point to sub-functionalization and antagonistic roles of RPA complexes in *C. elegans*.

## Introduction

Replication protein A (RPA) is a heterotrimeric complex which binds single-stranded DNA (ssDNA) with high affinity (reviewed in (1). In most organisms, this complex consists of a large (70 kDa), medium (32 kDa), and small (14 kDa) subunit (RPA1, RPA2, and RPA3 respectively in humans), each of which contains at least one oligosaccharide binding domain (OB fold), which gives the complex its ssDNA binding activity (1). RPA removes secondary structures in ssDNA, a property which is critical for replication and recombination (2). RPA was originally isolated as a factor that was essential for human simian virus SV40 *in vivo* replication (3). The role of RPA in replication is not only driven by its ability to bind ssDNA, but also through indirect interaction with proteins that are part of the replication machinery, including PCNA (4) and pol α (5). RPA also plays a role in cell cycle signaling and the DNA damage response, where RPA promotes ATM activation, possibly through its interaction with the MRN complex (reviewed in (6)), and ATR activation (7). In humans, the DNA damage induced apoptotic response is stimulated by RPA2 hyperphosphorylation (8). Furthermore, double-strand DNA break repair by homologous recombination (HR) also requires RPA, which involves its ssDNA binding property that is required for the assembly of the Rad51-ssDNA filament (reviewed in (9)). RPA is also required for other forms of DNA repair where ssDNA is formed (10). This complex is found in all eukaryotes, and the properties of the RPA complex appear to be conserved.

Not all organisms contain only the three canonical RPA subunits, and in some organisms, paralogs are found. Subunit paralog identities vary between organisms, and is driven by gene duplication events throughout evolution (11). The paralogs studied frequently retain the ancestral activities of the RPA subunit or lose some activities, but seldom neofunctionalise. For example, an RPA2 paralog, RPA4 is found in several mammals. In humans, RPA4 shares some activities with RPA2, where both facilitate homologous recombination. However, RPA4 is unable to signal cell-cycle progression or support replication (12). Plants have multiple copies of RPA1, RPA2, and RPA3 subunits, an outcome of their evolutionary history that involves many genome duplications. For example, the *Arabidopsis* genome contains five RPA1-like subunits, two RPA2-like subunits, and two RPA3-like subunits (13). The different RPA1 paralogs in *Arabidopsis* diverged in their functions: atRPA1C promotes meiotic HR, whereas atRPA1B and atRPA1D act in DNA replication. Archaea have RPA compositions that differ from eukaryotes, where some are missing RPA3 and only possess a large RPA1-like subunit, and one example has an RPA1-like subunit which dimerizes (14–16).

Gamete formation requires the faithful execution of two main functions supported by RPA: replication and recombination. Germ cells replicate their genome and undergo mitotic divisions in their stem cell niche to produce cells that enter meiosis. These cells are then required to repair a multitude of programmed DSBs by the process of recombination to produce the crossovers required for the formation of viable gametes. Crossovers act as a physical tether between homologous chromosomes, allowing for proper segregation of these chromosomes at the end of meiosis I. In many organisms, the absence of germline DSBs, or meiotic HR leads to the formation of egg and sperm that are inviable (17–20). In meiotic prophase I, DSBs form by the activity of the topoisomerase VI-like protein Spo11 (reviewed in (21)). Spo11 breaks are resected by nucleases in an MRN(X) dependent manner, leading to formation of ssDNA bound by RPA. To allow for strand invasion that leads to the formation of a double-Holiday junction, RPA is displaced by RAD51 (Reviewed in (22)). In the absence of RPA, RAD51 cannot efficiently assemble a RAD51-ssDNA filament leading to an inability to repair DSBs via HR. When HR is impaired, DSBs can be repaired through other mechanisms, such as canonical non-homologous end-joining (cNHEJ) and alternative end-joining (alt-EJ) (reviewed in (23)). In these repair events, ends are ligated together, often in an error-prone way, however, they do not promote proper meiotic chromosome segregation.

In *C. elegans*, there are orthologs for RPA1 (RPA-1) and RPA2 (RPA-2), as well as an additional subunit named RPA-4 which shares size and domain structure with RPA-2. Previously it was shown that RPA-1 forms foci in pachytene nuclei (24–27). RPA-1 foci are formed concurrently with RAD-51 focus formation, but continue to accumulate when RAD-51 foci numbers start to diminish (27). These finding suggest that RPA-1 serves two roles in recombination, one essential for RAD-51 loading on to ssDNA and one in localization at crossover intermediates (27). RPA-1 localization increases following treatment with DNA damaging agents such as hydroxyurea, UV, and ionizing radiation, likely due to accumulation of ssDNA in these cells and the requirement for RPA in HR and the replication stress response (25, 27) (28, 29). In a few studies, RPA-1 was shown to form a haze in mitotic germline nuclei(28). Knockdown of *rpa-1* by RNAi leads to embryonic lethality and defects in germline development (29, 30). These studies are consistent with an essential role for RPA-1 in DNA replication. While RPA-1 has been thoroughly studied, the additional subunits of RPA found in *C. elegans* have not, raising the question whether all RPA subunits serve identical functions.

Most germ cells of the *C. elegans* hermaphrodite germline undergo apoptosis leading to elimination of nuclei at the pachytene/diplotene transition. There are two known processes leading to germline apoptosis, one that is induced by stress (such as exogenously induced DNA damage), and one that occurs as part of germline development termed “physiological apoptosis” (31). Stress-induced apoptosis is considered to be a mechanism of removing nuclei that have selective disadvantage, such as those that contain excessive or unrepairable DNA damage. It is generally accepted that physiological apoptosis is required for germline nuclei to act as nurse cells/nuclei, contributing their cellular content to the oocytes that escape apoptosis (32). Mitogen-activated protein kinase (MAPK) signaling is required for licensing nuclei for apoptosis (32–34). MAPK acts on the core apoptotic machinery leading to activation of CEll Death abnormal-3 (CED-3, caspase) and CED-4 (Apaf-1), resulting in DNA fragmentation and the formation of a cell corpse (31). Cell corpses are then engulfed by CED-1 expressing somatic sheath cells to eliminate them from the germline (35). Germlines that are defective for apoptosis, produce abnormal and smaller eggs (36). A more extreme phenotype is observed when MAPK signaling is blocked, leading to pachytene-like nuclei accumulation in the proximal gonad (37). In *C. elegans,* RPA was not shown to play a direct role in apoptosis, however, RPA is involved in DNA damage signaling through the recruitment of ATR (ATL-1), and it may play a role in apoptosis signaling through its interaction with ssDNA (25).

Here we determine the roles of RPA-2 and RPA-4 in *C. elegans* meiosis. Our results show that RPA-1 and RPA-2 are involved in germline DNA replication, while RPA-4 is not. RPA-2 and RPA-1 are also critical for DSB repair, where these proteins localize to programmed and exogenously induced DSBs, while RPA-4 plays a more minor role in DSB repair. In contrast to other species, *C. elegans* RPA-2 is not completely required for RPA-1 focus formation nor is it essential for somatic replication. Surprisingly, RPA-4 localizes to DSBs formed by replication defects and exogenous damage, and attenuates the loading of RAD-51. Additionally, *rpa-2; rpa-4* double mutants display an unusual germline progression defect, and we demonstrate that this phenotype is due to defects in apoptosis. Altogether our studies point to unique functions of RPA in *C. elegans*.

## Results

### RPA-2, but not RPA-4, is required for mitotic replications

The *C. elegans* genome encodes for three RPA proteins (RPA-1, RPA-2, and RPA-4) identified by homology to RPA subunits from other organisms (wormbase.org, Figure 1A, Figure S1). RNAi of *rpa-1* results in gonad developmental defects (30) and RNAi of *rpa-2* results in sensitivity to radiation and DNA damage inducing agents (38). It is possible that the phenotypes observed by knockdown of genes encoding the RPA subunits is not fully representing the biological requirement for these genes, as it was performed by RNAi. To identify the roles of RPA subunits in *C. elegans* we created CRISPR/Cas9 mediated loss-of-function alleles for *rpa-1* and *rpa-4* and utilized an available deletion mutant for *rpa-2* (Figure S1). Consistent with its essential role in replication, an early frameshift mutant of *rpa-1* is larval lethal. Mutants with knockouts of genes essential only for HR in *C. elegans* develop due to maternal contribution to the zygote, but they lay dead eggs. However, *rpa-1* mutants of heterozygote mothers did not develop past the L2 larval stage and were diminished in their numbers (7% vs. the expected 25 % of progeny, Figure 1B). These observations are expected if RPA-1 is required for replication past the 100 cell stage (when maternal contribution is depleted), and if RPA-1 is an obligatory RPA subunit in replication. On the contrary, *rpa-2(ok1627)*, *rpa-4* single mutants and *rpa-2(ok1627)*; *rpa-4* double mutants develop to adults, indicating that these genes are not essential for somatic DNA replication. This result was unexpected, as in other metazoans RPA2 and RPA1 subunits are both essential for replication.

**Figure 1:**
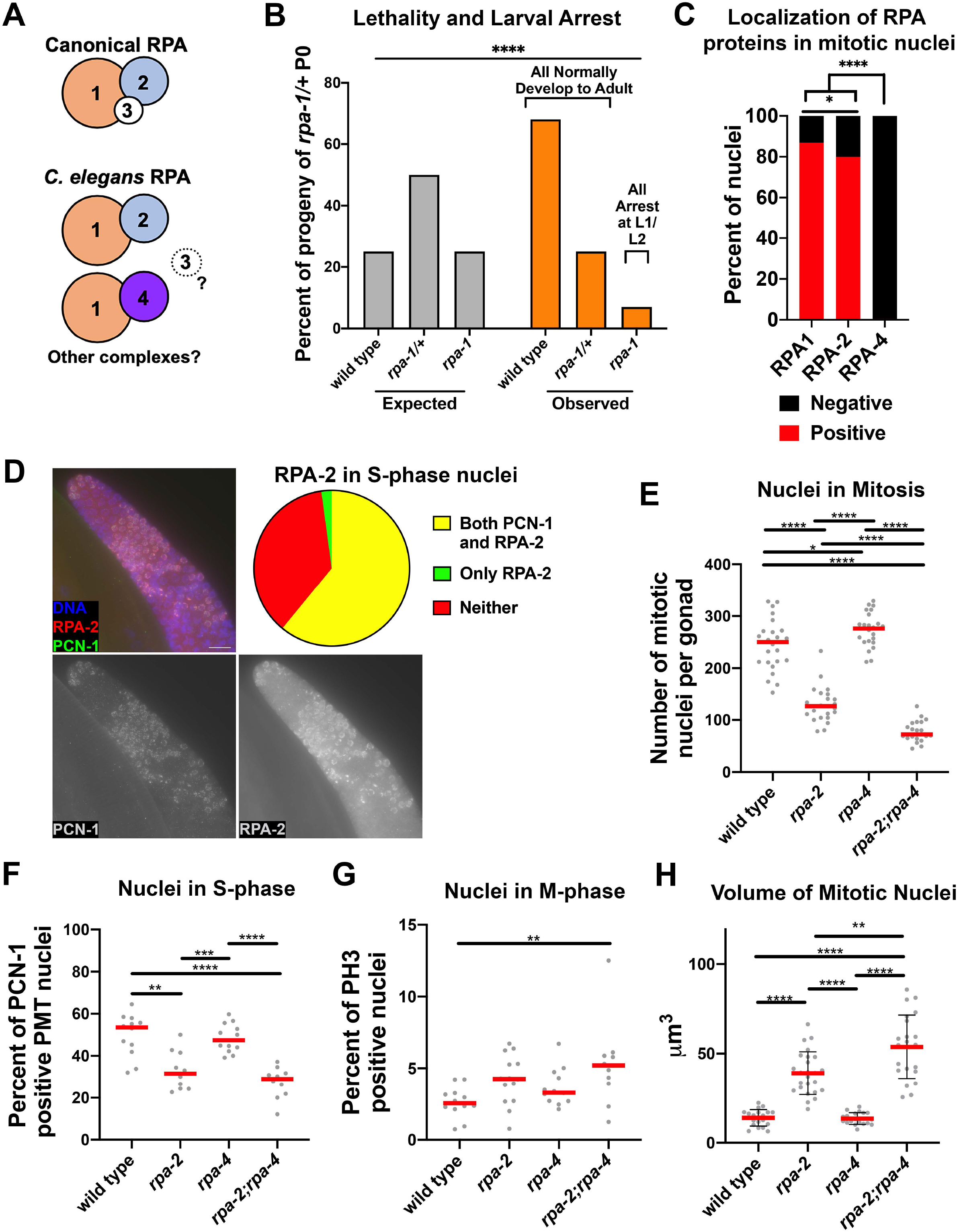
RPA-1 and RPA-2 are involved in replication in the pre-meiotic tip, where mutants of *rpa-2* and not *rpa-4* results in replication defects. A) Model for possible RPA complex combinations in *C. elegans.* RPA-4 and RPA-2 are interchangeable components of the complex, and RPA-3 is unknown. B) Larval lethality and arrest in rpa-1 mutants (Chi-square test performed, p-value<0.0001). C) Proportion of nuclei in the PMT with RPA staining for each of the tagged subunits. D) Co-staining of gonads with FLAG::RPA-2 and PCN-1. E) Number of mitotic nuclei as counted by position and morphology. F) Percent of nuclei with PCN-1 staining representing S-phase nuclei. G) percent of pH3 positive nuclei representing mitotic index. H) Nuclear volumes as estimated by calculations using FIJI acquired data (see materials and methods). Mann-whitney tests performed for C, E, F, and G, and T-test performed with welches correction for H, where p-values are represented as ****=<0.0001, ***=<0.001, **=<0.01, and * =<0.05)

To determine the function of RPA subunits in *C. elegans* meiosis, we N-terminally tagged *rpa-2* and *rpa-4* using CRISPR with 3X FLAG epitopes. RPA-1 was previously N-terminally tagged using the OLLAS epitope, but the localization pattern was not described in detail (28). All epitope tagged proteins are likely functional, as epitope tagged strains showed no difference in brood size or egg viability ((28)and Figure S2A). The RPA complex has a well-established role in replication (39), therefore we examined the localization of the three RPA subunits in nuclei undergoing replication in the germline. The pre-meiotic tip (PMT) contains the stem cell niche of the germline and most nuclei are found in S/G2 phases of the cell cycle (40, 41). OLLAS::RPA-1 and FLAG::RPA-2 staining was present in a majority (87% and 80% respectively) of nuclei of the PMT (Figure 1C), which is similar to the previously reported percent of S-phase nuclei (~60-70% (40, 41)). Most of these nuclei form abundant OLLAS::RPA-1 and FLAG::RPA-2 foci co-localizing with chromosomes (DAPI) which appear as a “haze” or “coating” on the DNA. To determine if this RPA localization pattern is associated with S-phase nuclei, we co-stained for FLAG::RPA-2 and PCN-1 (PCNA homolog, clamp subunit associated with DNA polymerase during replication, (42, 43)). PCN-1 is found in S-phase nuclei in a similar localization pattern to that of RPA-1 and RPA-2, and all nuclei that expressed PCN-1 staining also stained for FLAG::RPA-2 (Figure 1D). *rpa-4* appears to be a complex duplication with several isoforms, the largest product of which is isoform a, which shares 30.56% identity to RPA-2. RPA-2 and RPA-4 appear to be orthologs of human RPA-2 based on their protein alignment (44). Therefore, we suspected that the two proteins would exhibit a similar localization pattern. However, unlike RPA-2, FLAG::RPA-4 was almost completely absent from PMT nuclei (Figure 1C) suggesting that RPA-4 does not play a role in replication.

Next we examined the role of RPA subunits in germline DNA replication through the analysis of germlines in *rpa-2* and *rpa-4* mutants. Since no worm past the L2 stages were observed in *rpa-1* mutants, we were unable to use this allele to further study RPA-1’s role in the germline, because it is tissue that develops in later larval stages. *rpa-2* and *rpa-4* single mutants as well as *rpa-2*; *rpa-4* double mutants all contained germlines, indicating that neither RPA-2 or RPA-4 is required for replication leading to germline formation. However, detailed analysis of *rpa-2* mutants uncovered a role for RPA-2 in germline DNA replication. We compared the total number of mitotic nuclei in the PMT in *rpa-2* knockout worms based on nuclear morphology. In *rpa-2(ok1627)* young-adult hermaphrodites, the total number of mitotic nuclei was almost half of the amount that is observed in wild-type germlines (Figure 1E). We tested if this reduction could be attributed to a decrease in the number of S-phase nuclei by quantifying the number of nuclei with PCN-1 staining. The percent of S-phase nuclei in *rpa-2* single mutants and *rpa-2(ok1627); rpa-4(iow24*) double mutants was reduced by nearly half, indicating a reduction in S-phase nuclei (Figure 1F). Next, we stained and quantified the number of nuclei that were positive for phosphorylated histone H3 (PH3), which marks mitotic M-phase nuclei (Figure 1G). We observed a small increase in the percent of M-phase nuclei, consistent with M-phase arrest triggered by replication errors. Germline nuclei that experience replication block, such as those that are treated with hydroxyurea (HU), arrest and increase their nuclear volume (25). In concordance with the presence of replication defects, *rpa-2* mutants had larger nuclei than wild type (Figure 1H and S2B). If replication defects lead to decreased mitotic nuclei, it is likely that this could lead to reduction in overall germline size, as less nuclei enter meiosis. Indeed, gonad length was reduced in *rpa-2* mutants (Figure S2C). All together, these results indicate that RPA-2 is essential for normal germline replication.

Since RPA-4 shares homology with RPA-2, it is possible that RPA-4 functions similarly to RPA-2, such that *rpa-4(iow21)* mutants will phenocopy *rpa-2* mutants. However, RPA-4 localized in few mitotic nuclei (Figure 1C), suggesting it may not play a role in mitotic germline nuclei. *rpa-4(iow21*) mutants where indistinguishable from wild type in the percent of PCN-1 positive nuclei and only showed mild effects on total nuclei numbers or PH3 positive nuclei, supporting our hypothesis that RPA-4 does not play a significant role in replication. In agreement, *rpa-4* mutants did not modify the *rpa-2* phenotype in the parameters indicative of effect on replication, and *rpa-2(ok1627); rpa-4(iow24)* double mutants only had small effects on total mitotic nuclei number and nuclear volume compared to the *rpa-2(ok1627)* single mutant (PCN-1 or PH3 positive nuclei, figure 1D and 1G). *rpa-2(ok1627); rpa-4(iow24)* mutants had longer gonads than *rpa-2(ok1627)* mutants, which we discuss below may be due to a later meiotic role of RPA-4. These data are consistent with limited localization of RPA-4 in mitotic nuclei, suggesting that despite its homology to RPA-2, RPA-4 does not play a significant role in replication.

### RPA-1 and RPA-2 colocalize and form a complex that excludes RPA-4 in wild type pachytene nuclei

During meiotic prophase I, programmed DSB are formed by SPO-11 and repaired by HR (20), which requires the RPA complex for proper RAD-51 loading (24). RPA-1 was previously shown to localize to germline nuclei, however the localization of RPA-2 has not been reported. To identify the localization pattern of RPA-2 in meiotic cells undergoing recombination we stained gonads of our 3XFLAG tagged RPA-2 strain, and compared the localization to that of OLLAS tagged RPA-1. The gonads were divided into 7 zones (for details see materials and methods), where zones 1-2 represent mostly the mitotic zones, zone 3 represents mostly the transition zone (Leptotene and Zygotene), and zones 4-7 represent pachytene. RPA-1 and RPA-2 have similar localization patterns in the *C. elegans* hermaphrodite germline (Figure 2A and 2B). RPA-1 and RPA-2 are mostly absent from the transition zone (Z3), where programmed meiotic DSBs form (24). During pachytene, DSBs are resected and RPA binds the ssDNA ends. Interestingly RPA-1 and RPA-2 foci numbers increase from early to late pachytene, which was not expected as RAD-51 (which displaces RPA) focus formation peaks at early pachytene (Figure 6A). However, these results match previous observations of RPA-1 foci in late pachytene in nuclear spreads, which likely mark cross-over intermediates (27). We observed in late pachytene/diplotene nuclei a “haze” of RPA-1 and RPA-2 similar to what was observed in the PMT, which we interpret as nuclei preparing for the next round of replication after fertilization. In the pachytene region, RPA-2 foci were less abundant than RPA-1 (1 focus/nucleus compared to 4.4 foci/nucleus, respectively), suggesting RPA-1 can bind ssDNA without RPA-2.

**Figure 2:**
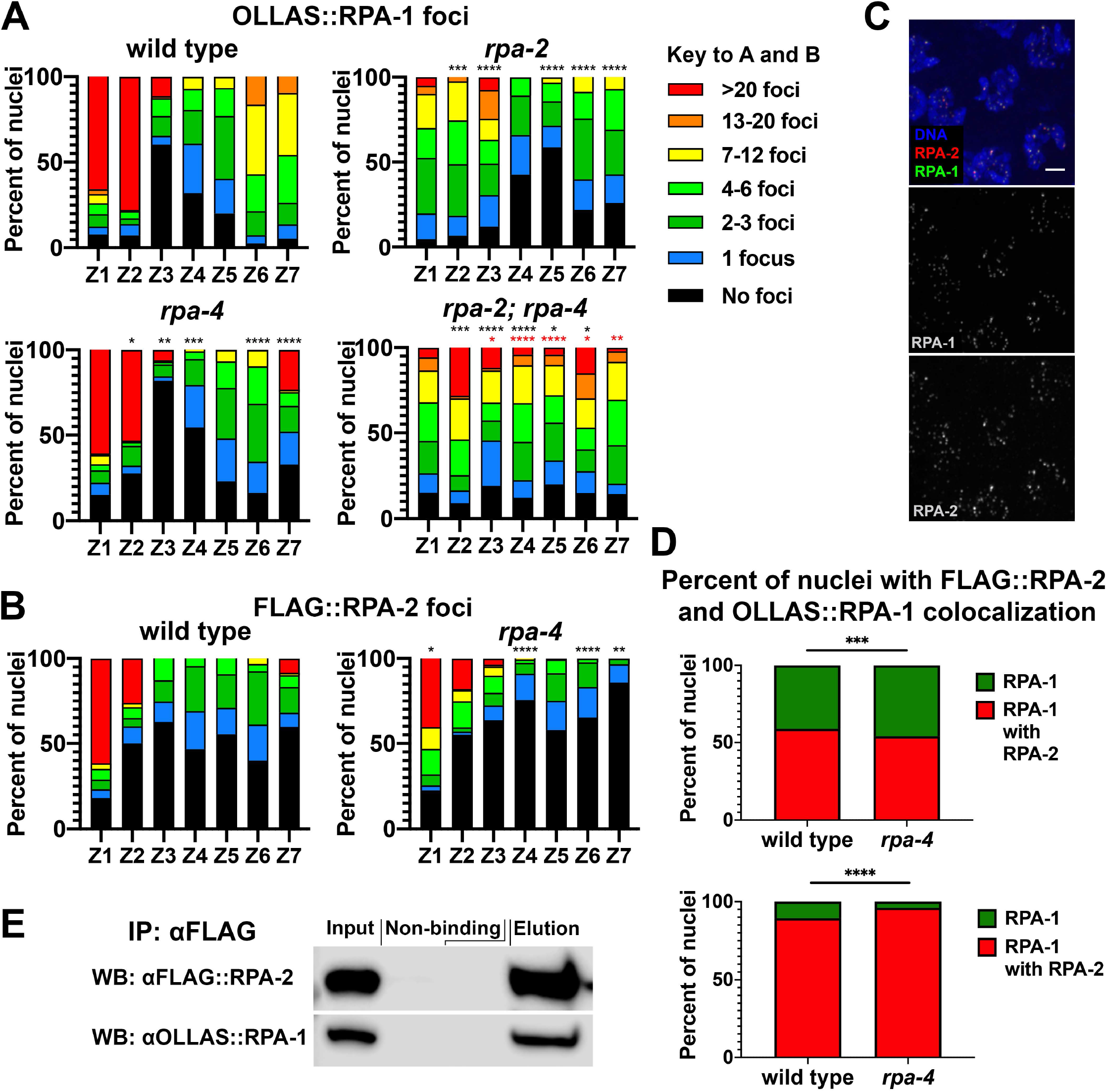
RPA-1 and RPA-2 colocalize and interact *in vitro*; interaction is at DSBs in pachytene. A) Percent of nuclei with indicated amount of OLLAS::RPA-1 foci. Black asterisks are comparison with wild type, and red asterisks are comparison with *rpa-2* mutants. B) Percent of nuclei with indicated amount of FLAG::RPA-2 foci. (Black asterisks represent comparison with wild-type, and red asterisks represent comparison with *rpa-2*) *C*) Image of mid-pachytene nuclei showing colocalization of FLAG::RPA-2 and OLLAS:RPA-1 in otherwise wild-type background. D) Percent of colocalization of FLAG::RPA-2 and OLLAS:RPA-1 in pachytene zones 4-6. E) Pull down of FLAG::RPA-2 with Co-IP of OLLAS::RPA-1. (Mann-Whitney tests performed, where p-values are represented as ****=<0.0001, ***=<0.001, **=<0.01, and * =<0.05)

In most species, RPA-1 and RPA-2 form a complex required for solubility and stability of RPA-1, and thus RPA-2 is essential for RPA-1 function, including its binding to DSBs (45). However, deletion of *rpa-2* reduced, but did not eliminate OLLAS::RPA-1 focus formation (Figure 2A). The most robust effect of *rpa-2* mutants on RPA-1 localization was in mitotic cells (zones 1 and 2), indicating a role for RPA-2 in promoting the interaction of RPA-1 with ssDNA for normal replication, as supported by PCN-1 staining (Figure 1D). *rpa-2* knockout also had an effect on OLLAS::RPA-1 loading in meiotic nuclei, where zones 5-7 have fewer OLLAS::RPA-1 foci than wild type (49%, 68%, and 62% reduction in each respective zone, Figure 2A), indicating that RPA-2 is also required for wild-type levels of RPA-1 localization during DSB repair.

Next we examined if RPA-1 and RPA-2 co-localize *in vivo* at the time of meiotic DSB repair. FLAG::RPA-2 and OLLAS::RPA-1 foci co-localize extensively, consistent with formation of an RPA complex including these two subunits (Figure 2D). Almost all RPA-2 foci colocalized with RPA-1, but not all RPA-1 foci colocalized with RPA-2, consistent with higher abundance of RPA-1 foci compared to RPA-2 foci. Co-immunoprecipitation (co-IP) supports that this co-localization reflects a physical interaction between RPA-2 and RPA-1, as pull-down of FLAG::RPA-2 resulted in co-IP of OLLAS::RPA-1 (Figure 2E). Unlike *rpa-2* deletions, *rpa-4* deletion only had a minor effect on the localization of RPA-1 when compared to wild-type gonads. Co-localization of RPA-1 with RPA-2 foci was slightly reduced in the *rpa-4* mutant backgrounds, suggesting that RPA-4 does little to affect the interaction of RPA-2 with RPA-1 (Figure 2D).

### RPA-4 localizes to subset of DSBs, regulated by RPA-2 and inhibits RPA-1 focus formation

RPA-4 is the ortholog of RPA-2, suggesting it may have a function in HR. Similar to mitotic germline nuclei, RPA-4 was also absent from most meiotic germline nuclei. Overall, FLAG::RPA-4 was found in less than 1% of germline nuclei, and almost exclusively in the pachytene stage of meiotic prophase I (figure 3A). The localization of RPA-4 in the form of foci in meiotic nuclei suggests that it is recruited to the sites of DNA damage, likely ssDNA. The limited localization of RPA-4 to meiotic nuclei suggests that RPA-4 localized only to a subset of DSBs. Germline DSBs are generated by 3 mechanisms: SPO-11 induced DSBs, DNA damage due to replication fork collapse and DNA fragmentation following apoptosis. SPO-11 induced DSBs are the majority of germline DSBs and are found only in early to mid-prophase. DNA fragmentation following apoptosis is only found in few nuclei in the late pachytene region of the germline. The localization of RPA-4 to nuclei prior to late pachytene indicates that it is not marking fragmented DNA of apoptotic nuclei (but this doesn’t preclude it from binding DNA of nuclei marked for apoptosis). To test if RPA-4 localization depends on programmed meiotic DSBs, we tested FLAG::RPA-4 is *spo-11* mutants. In agreement with its limited localization, RPA-4 foci numbers were not affected by the removal of *spo-11*, indicating that RPA-4 marks SPO-11-independent DSBs (Figure 3A). Therefore, it is likely that under normal growth conditions, RPA-4 marks DSBs created by other forms of DNA damage such as replication fork collapse. This DNA damage may not be repaired in mitotic nuclei and is carried over as these nuclei proceed to meiosis.

**Figure 3:**
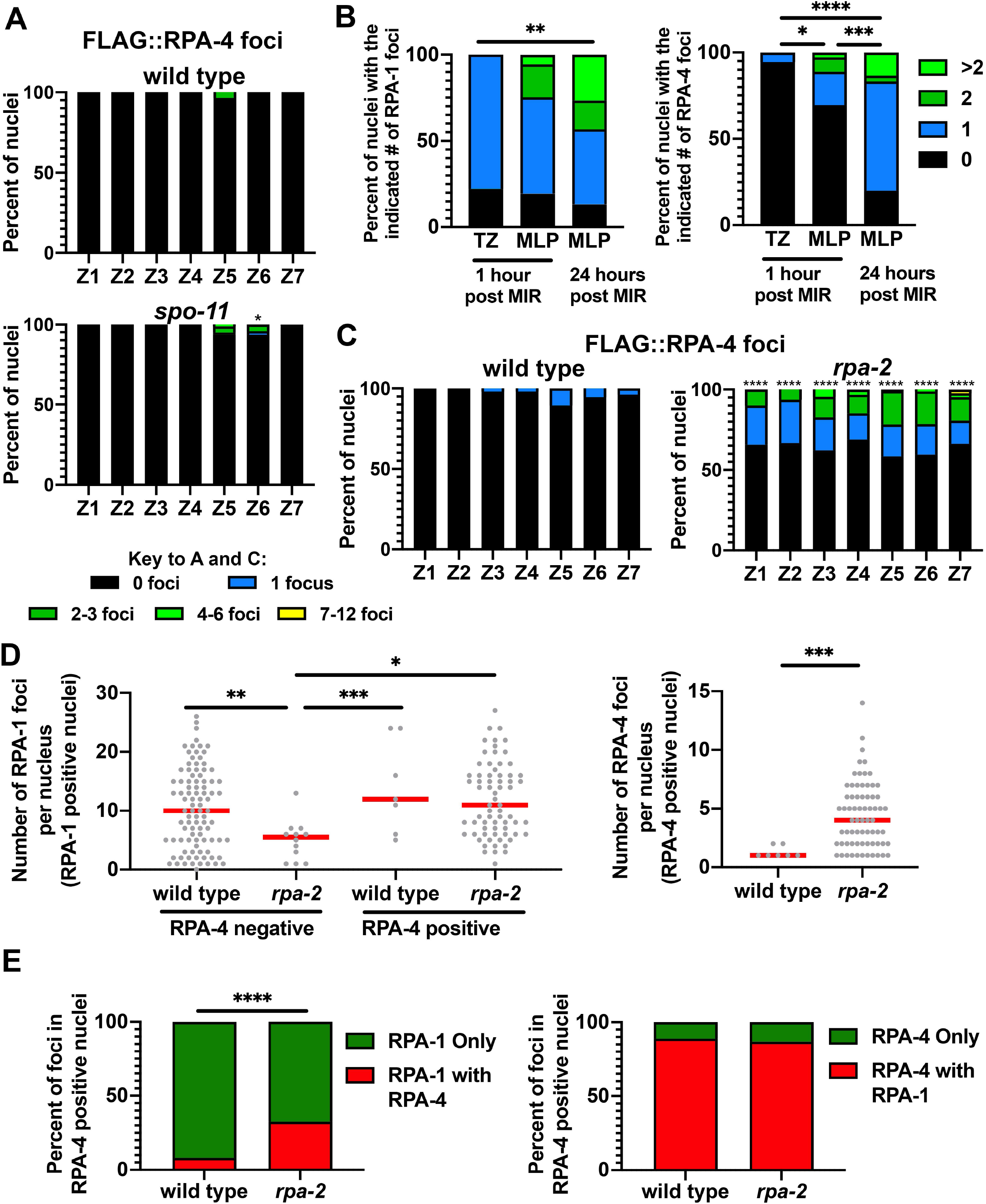
RPA-4 forms rare foci in wild-type worms that become more abundant in *rpa-2* mutants, and localize to SPO-11-independent and exogenous DSBs following RPA-1. A) Percent of FLAG::RPA-4 foci in *spo-11* mutants. B) Abundance of OLLAS::RPA-1 and FLAG::RPA-4 MIR foci in indicated zones and time periods. C) Percent of nuclei with indicated number of FLAG::RPA-4 foci in wild type and *rpa-2* mutant gonads. Percent Colocalization of FLAG::RPA-4 foci with OLLAS::RPA-1 MIR foci (TZ= transition zone (Leptotene/zygotene), MLP = Mid-to-late-pachytene). D) number of RPA-1 and RPA-4 foci in RPA-4 positive and negative mid-pachytene nuclei. E) Percent of FLAG::RPA-4 and OLLAS::RPA-1 foci in mid-pachytene nuclei that colocalize. (Mann-Whitney tests performed, where p-values are represented as ****=<0.0001, ***=<0.001, **=<0.01, and * =<0.05)

To test if RPA-4 focus formation is dependent on DSBs, we performed UV laser micro-irradiation (MIR) in adult hermaphrodite germlines. For this study, we used GFP11 tagged RPA-1 expressed from the endogenous locus to mark DSB sites [previously generated in our lab: (28)]. We found that indeed, GFP11 tagged RPA-1 localized to the site of exogenously induced DSBs forming about 1 focus per nucleus and localized to DSBs 8-10 minutes following MIR (Figure S3A). These findings are consistent with ones observed with a different transgenic line (integrated extra-chromosomal array (46)). Using an OLLAS tagged RPA-1, we observed RPA-1 localization that increased 24 hours post MIR (Figure 3B. RPA-1 MIR foci are detected in higher levels in mid-pachytene nuclei compared to transition zone nuclei (Figure 3B), consistent with previously published data (46)). To test if RPA-4 is recruited to exogenously induced DSBs we analyzed the co-localization of RPA-4 with RPA-1 following laser MIR. Our results indicate that all (100%, n=54) RPA-4 localizes to MIR foci containing RPA-1. Unlike RPA-1 which localized to MIR damage in the first hour post MIR, RPA-4 foci were more apparent only 24 hours following MIR (Figure 3B). Taken together, these data suggest that RPA-4 foci localize to DSBs and that it follows RPA-1 localization. RPA-4 localization likely depends on resection, and likely forms a complex with RPA-1.

Given the distinct localization pattern of RPA-2 and its paralog RPA-4, it is possible that the two proteins have different functions. While *rpa-4* had only minor effects on RPA-2 localization, *rpa-2* removal showed a notable effect on RPA-4 localization. In *rpa-2* worms, the number RPA-4 positive nuclei increased by an average of 4-fold compared to wild type (from 8 foci/gonad in wild-type worms to 128 foci/gonad in *rpa-2* worms, Figure 3C), indicating that RPA-2 attenuates RPA-4 focus formation, either directly or indirectly (see discussion). The increase in RPA-4 focus formation in *rpa-2* mutants is likely due to the increase in DNA damage due to replication defects (Figure 1).

To investigate the relationship between RPA-4 and RPA-1, we focused our attention on RPA-4 positive nuclei (Figure 3D). RPA-4 foci were only found in nuclei that also contained RPA-1 foci. RPA-4 positive nuclei had similar levels of RPA-1 foci when compared to RPA-4 negative nuclei (14 vs 10.5 foci/nucleus in pachytene) in wild type worms. However, in *rpa-2* mutants, RPA-1 foci were more abundant in RPA-4 positive nuclei, as compared to RPA-4 negative nuclei (11.8 vs 5 foci/nucleus in pachytene (Figure 3D)). As evident by the overall distribution of RPA-4 foci (3C), when focusing only on RPA-4 positive cells, RPA-4 levels were significantly elevated in the absence of *rpa-2*. While RPA-1 were more abundant that RPA-4 foci, in RPA-4 positive cells, almost all RPA-4 foci colocalized with RPA-1 in wild type and *rpa-2* mutants (Figure 3E). Altogether these data suggest that RPA-4 focus formation is down regulated by RPA-2, dependent upon significant amounts of unrepaired DNA damage.

### *C. elegans* RPA-1, RPA-2 and RPA-4 form RPA-1/RPA-2, RPA-1/RPA-4 and RPA-2/RPA-4 complexes

Using co-IPs, we have shown that RPA-1 and RPA-2 physically interact. RPA-4 was not detectable on Western blot analysis due to its low abundance which prevented the analysis of its physical interaction with RPA-2 and RPA-1 by traditional co-IP. To bypass this limitation, we performed Single-Molecule Pull Down (47) experiments using a triple tagged strain containing OLLAS::RPA-1, 3XFLAG::RPA-2 and MYC::RPA-4. The SiMPull experimental strategy involved capture of the RPA complexes from the whole worm lysate by the surface-tethered biotinylated antibodies against a tag on one of the subunits, followed by visualization of the RPA-1, RPA-2 and/or RPA-4 proteins via fluorescent antibodies specific to tags present on each protein. This approach allowed us to enumerate RPA complexes of different compositions present in the same mixture. We expected to see complex formation of RPA-1 with RPA-2 predominately as it was detected in IPs (Figure 2E) and because the recombinant RPA-1 and RPA-2, when co-expressed, have been shown to form a 1:1 heterodimeric complex (48). Unexpectedly, we found evidence for formation of three distinct complexes including RPA-1/RPA-2, RPA-1/RPA-4 and RPA-2/RPA-4 (Figure 4B-C). In a very few instances (about 1% of observed events), the RPA-1/RPA-2/RPA-4 complexes were also observed. The bar graphs in Figure 4 (B-C) show track counts taken across multiple fields of view for each respective combination. RPA complexes containing OLLAS::RPA-1 were captured from the lysate to the surface-tethered anti-OLLAS antibodies (Figure 4C). The presence of 3XFLAG::RPA-2 and MYC::RPA-4 was simultaneously detected and quantified using Cy3-labeled anti-FLAG and Cy5-labeled anti-MYC antibodies, respectively. Pre-Ab track and wild-type control track counts represent the number of counts before the fluorescent antibodies are added into the microscope flow cell and the control experiment that uses the lysate from the wild type animals, respectively. Post-Wash values represent the number of tracks measured after the antibody had incubated for 30 minutes in the presence of lysate and had been washed with imaging buffer, which represents the number of respective RPA complexes. Unexpectedly, we observed RPA-1/RPA-2 and RPA-1/RPA-4 with the same frequency. Then, 3XFLAG::RPA-2 was anchored to the surface and OLLAS::RPA-1 and MYC::RPA-4 were visualized using Cy3 and Cy5-labeled antibodies against OLLAS and MYC, respectively (Figure 4CD). Next, we anchored the MYC::RPA-4 and visualized RPA complex formation by adding fluorescent antibodies against OLLAS (RPA-1) and FLAG (RPA-2), which recapitulated our previous results (Figure 4D and E). In both configurations, we observed RPA-4 preferentially binding RPA-2. There was no RPA-1/RPA-2/RPA-4 complex formation in the Pre-Ab and wild type controls (colocalization of Cy3 and Cy5 dyes), however three events were noted in the Post-Wash values in all experiments. While unlikely, the appearance of these rare complexes may represent an artifact of the method as it only accounts for less than 1% of the total counts. These complexes may arise from two biotinylated antibodies located within diffraction limited spot on the surface (closer than 250 nm). These signals, however, may also represent actual RPAs whose functions need to be further investigated. Combining this analysis with the analysis performed above we conclude that RPA-4 acts independently of RPA-2 in regulating RPA-1 activity, which may involve RPA-2/4 interaction (Figure 4D).

**Figure 4:**
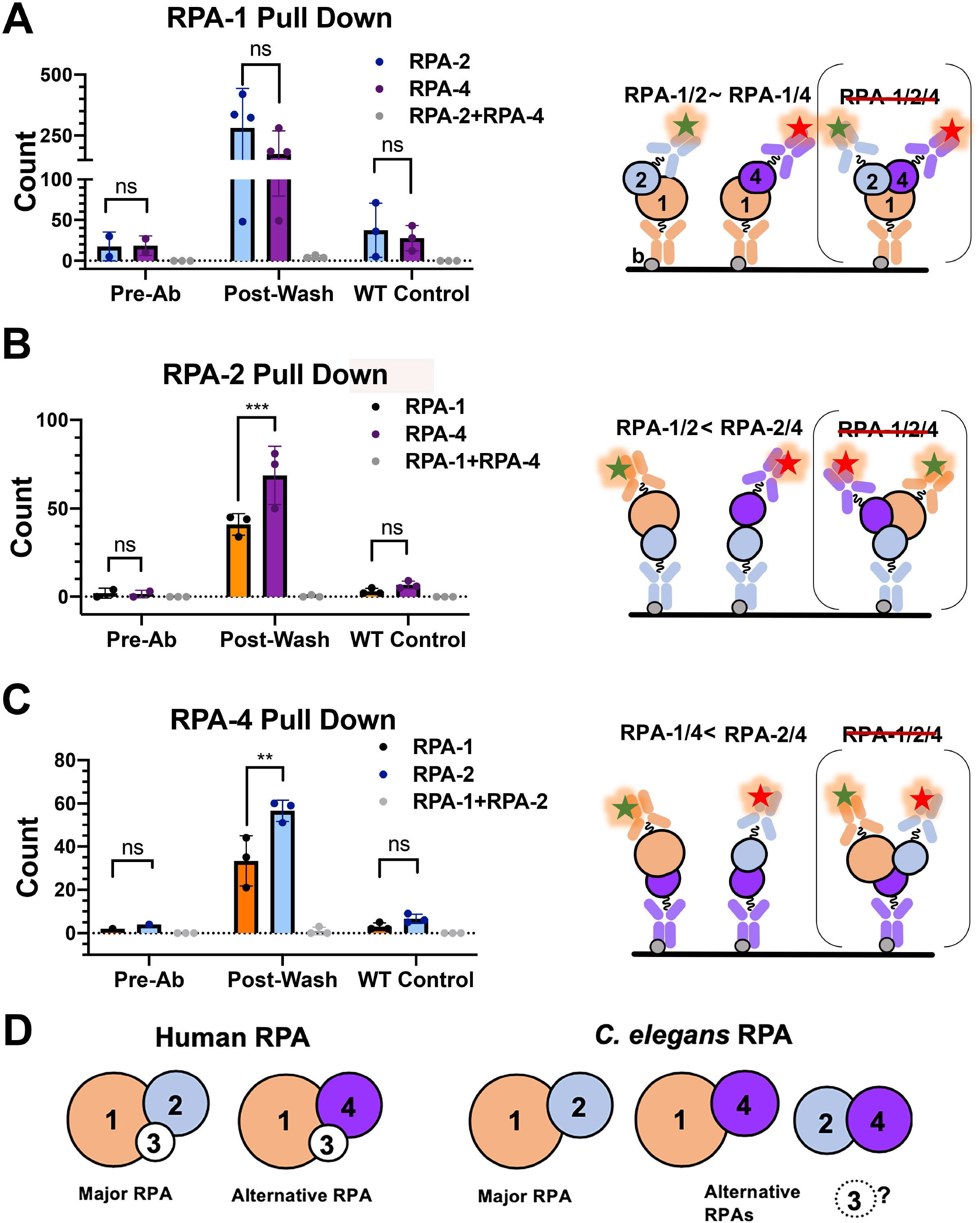
Single-molecule pulldown reveals presence of 3 possible complex arrangements. A) Number of RPA-4 and RPA-1 pull down counts through interaction with RPA-2. B) Number of RPA-2 and RPA-4 pull down counts through interaction with RPA-4. C) Number of RPA-2 and RPA-4 pull down counts through interaction with RPA-1. The data are shown for three independent experiments. Pre-Ab is a control for non-specific signals in the flow cell treated with respective biotinylated antibody and worm extract. Wild type (WT) control reflects the experiments using worms expressing un-tagged RPA forms. The data for each pulled pair were compared using two-way ANOVA. The respective p value is shown under the graph. D) Model interpretation of the results in comparison to human RPA complex.

### RPA-2, but not RPA-4, is essential for meiotic recombination

Proper repair of meiotic DSBs is critical to the viability of the resulting gametes. If RPA-2 performs an essential role in meiotic DSB repair, we expect it to be essential for embryonic viability. As expected, deletion of *rpa-2* resulted in reduction in embryonic viability, where no eggs hatched in single mutants or *rpa-2; rpa-4* double mutants (Figure 5A). These data suggest that homologous repair of DSBs requires RPA-2. To address this possibility, we examined the diakinesis nucleus adjacent to the spermatheca (diakinesis-1). In wild type worms, we expect to observe six DAPI bodies in this nucleus, representing the six pairs of homologous chromosomes joined by crossover. Lack of DSB repair will result in DNA fragmentation as each chromosome has several DSBs (24). However, in *rpa-2* hermaphrodites, a range of different DAPI body numbers was observed compared to wild type, indicating that repair of meiotic DSBs occurred. The irregular size of these DAPI bodies indicates that repair did not use homology and likely involved error-prone DSB repair mechanisms (Figure 5B). To test this, we crossed to mutants of *cku-70* (part of the canonical non-homologous end joining (cNHEJ) pathway) and/or *polq-1*(part of the alternative endjoining pathway) in *rpa-2* mutants. While removing either one of these pathways by itself had no effect on the number of DAPI bodies, an increase was observed in the triple mutant *polq-1; cku-70; rpa-2* strain, confirming the repair of breaks occurs through canonical and alternative end-joining (error-prone) repair mechanisms, and that these pathways act redundantly to repair unrepaired DSBs in the germline (Figure S4A). Thus, while RPA-1 is essential for viability starting from early development (Figure 1B), RPA-2 is essential later in development, for reproduction.

**Figure 5:**
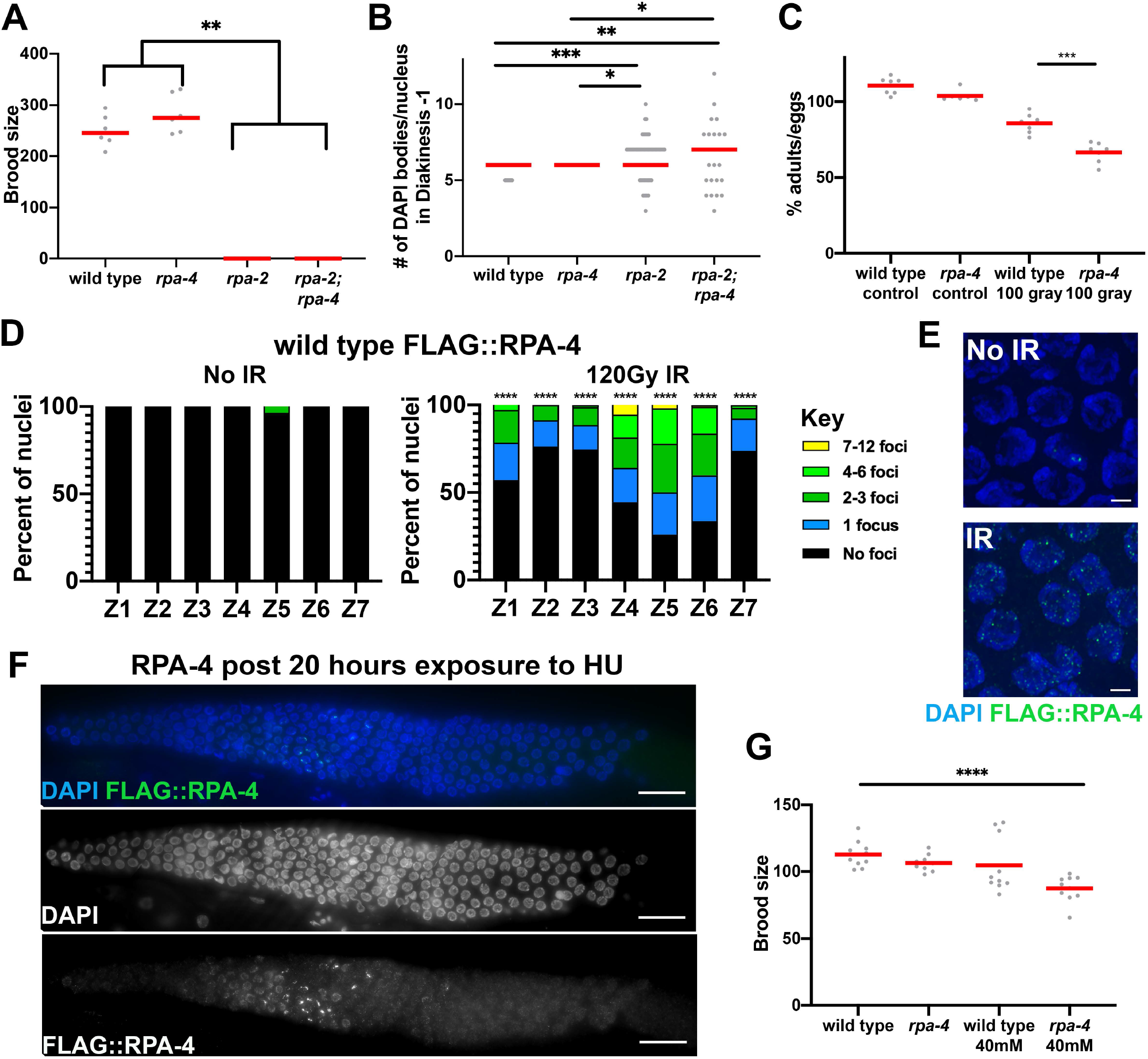
Deletion of *rpa-2* and not *rpa-4* leads to defects in meiotic HR, FLAG::RPA-4 localizes to exogenously induced DSBs, and *rpa-4* deletion leads to HU and IR sensitivity. A) Brood size for each mutant genotype, each point represents the number of adult progeny from a single parent. B) Number of DAPI bodies counted in diakinesis −1 nuclei. C) Percent viability of eggs laid in wild type and *rpa-4* mutants that have been irradiated with 100Gy gamma-IR and the respective controls. D) Percent of nuclei with the indicated number of FLAG::RPA-4 foci. E) Representative image of mid-pachytene nuclei with and without gamma-IR. F) Representative images of FLAG::RPA-4 gonad stained 20 hours after treatment with 40mM HU. G) Brood size of HU treated wild type and *rpa-4* mutant worms. (Mann-Whitney tests performed, where p-values are represented as ****=<0.0001, ***=<0.001, **=<0.01, and *=<0.05) E) Time of appearance for the first RAD-51 MIR focus (p-value=0.0847).

### RPA-4 localization is upregulated in response to induced DNA damage

Despite the similarity to RPA-2, RPA-4 may not be required under normal growth conditions, in which DSBs are programmed. To test whether it is required following exposure to exogenous DNA damage, we introduced DNA damage by ionizing radiation (IR). This DNA damage is thought to primarily induce DSBs and occurs throughout the germline. When analyzing FLAG::RPA-4 localization following exposure to gamma-IR, RPA-4 localization increased in abundance from <1% to about half of nuclei (Figure 5D and E). To test whether RPA-4 is recruited to other types of exogenous DNA damage, we exposed worms to hydroxyurea (HU), which creates DSBs by inducing replication stress. RPA-4 localized extensively to transition zone and early pachytene nuclei, with very few foci in mitotic zone nuclei, despite HU damage being induced at this stage (Figure 5F). RPA-4 presence in nuclei after exit from mitosis (after DNA damage was processed) is consistent with the observation that RPA-4 is not recruited to DSBs immediately and that it is preferentially localized to pachytene nuclei in unexposed germlines (Figure 3A). To test if the recruitment of RPA-4 to HU breaks effects viability, we tested for the brood size of *rpa-4* mutants following HU exposure. *rpa-4* mutant worms exposed to HU showed a reduction in their brood size following exposure to HU when compared to wild-type unexposed controls (Figure 5G). Altogether these data show that RPA-4 is recruited to DSBs when their levels are increased (*rpa-2* mutants, ionizing irradiation or HU exposure).

### RPA-4 inhibits RAD-51 focus formation

RPA is required for efficient RAD-51 recruitment to ssDNA, an essential step in the repair by HR (3, 49, 50). Therefore, if RPA loading is impaired, we would not expect RAD-51 to form foci in pachytene nuclei. *rpa-2* deletion severely reduces but did not eliminate the presence of RAD-51 foci, confirming a defect of meiotic HR as the source of the abnormal DAPI body phenotype (Figure 5D). Unlike *rpa-2* mutants, *rpa-4* mutants showed only mild effects under normal growth conditions. *rpa-4* mutant progeny had similar embryonic viability compared to wild type under normal growth conditions (Figure 5C). Next, we tested if RPA-4 is required for embryo viability when the germline is challenged with exogenous DNA damage. Following 100 Grays of ionizing radiation from a cesium source, viability of *rpa-4* mutant worms was significantly reduced compared to irradiated wild-type control worms, indicating a sensitivity to ionizing radiation in *rpa-4* worms (Figure 5C and Figure S4B). These data suggest that RPA-4 plays a significant but smaller role compared to RPA-2 in recombination.

Despite the seemingly wild-type localization of RAD-51 foci in *rpa-4* mutants, the timing of RAD-51 foci appearance was altered. We observed greater numbers of RAD-51 foci in zone 4 and a reduction of RAD-51 foci numbers in zones 5-7 in *rpa-4* mutants compared to wild type (Figure 6A). This may indicate more rapid RAD-51 loading and removal in the absence of *rpa-4*, which is consistent with a model by which RPA-4 inhibits RPA-1 recruitment. In agreement, *rpa-2; rpa-4* double mutants had significantly more RAD-51 foci than *rpa-2* single mutants (zone 1, and 5-7). To test this possibility, we analyzed the timing of recruitment of GFP::RAD-51 to MIR foci in the presence and absence of *rpa-4*. This analysis was performed as previously described in *spo-11* mutant background to reduce background levels of foci (46). As expected, UV laser microirradiation RAD-51::GFP foci were significantly more abundant, and loaded slightly faster (but not significantly) in *rpa-4* mutants (Figures 6B and C), indicating that RPA-4 indeed inhibits RAD-51 focus formation.

**Figure 6:**
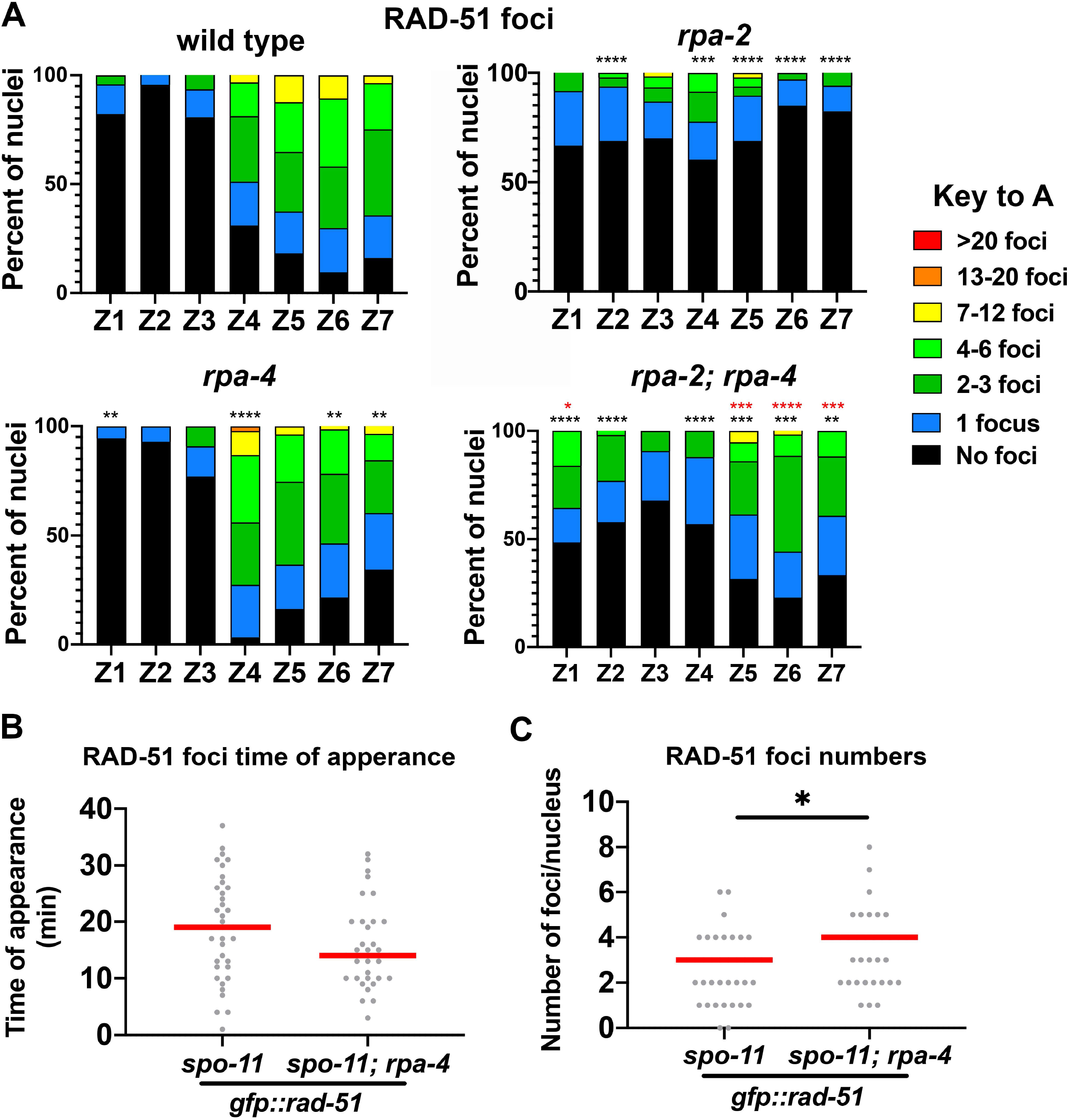
*rpa-2* and *rpa-2; rpa-4* mutants have decreased RAD-51 foci compared to wild type, and RAD-51 MIR foci are more abundant in *rpa-4* mutants. A) Percent of nuclei with indicated amount of RAD-51 foci. Black asterisks are comparison with wild type, and red asterisks are comparison with *rpa-2* mutants. (Mann-Whitney tests performed, where p-values are represented as ****=<0.0001, ***=<0.001, **=<0.01, and *=<0.05, where black asterisks represent comparison with wild-type, and red asterisks represent comparison with *rpa-2*) B) Time of appearance of GFP::RAD-51 foci in *spo-11* mutant background after treatment with UV Laser MIR. C) Number of GFP::RAD-51 foci following treatment with UV Laser MIR. (Mann-Whitney, p-value=0.0847).

### RPA-4 promotes germline apoptosis, in *rpa-2* mutants

In late pachytene about half of meiotic nuclei are eliminated by apoptosis while surviving nuclei and their chromosomes go through characteristic morphological changes as they transition to diplotene and then diakinesis. These changes are observed in the bend region of the germline, where the gonad that is positioned inside the worm’s body bends towards the uterus of the worm, about halfway through the length of each gonadal arm. *rpa-2* and *rpa-4* mutants exhibited normal progression into diakinesis as found in wild type (Figure 7A). Analysis of the *rpa-2; rpa-4* double mutants uncovered a surprising role for the paralogs in germline progression. Pachytene nuclei were observed extending past the bend in *rpa-2; rpa-4* germlines, where diplotene and diakinesis meiotic stages should occur (Figure 7A). The distance from the bend to the first diakinesis nucleus was slightly increased in *rpa-4* mutants compared to wild type or *rpa-4* mutants. However, *rpa-2; rpa-4* mutants exhibited a 2.5-fold increase in this measurement (~36μm compared to 214μm, figure 7B). Fewer diakinesis nuclei were observed in *rpa-2* mutants than wild type or *rpa-4* mutants (Figure 7C). This was an expected outcome of the reduction in germline proliferation in the *rpa-2* mutant background, leading to reduced numbers of mitotic germline nuclei, and germline length, as shown above. Despite identical effects on germline proliferation (Figure 1E-H), diakinesis nuclei were scarcely found in *rpa-2; rpa-4* double mutants, compared to *rpa-2* mutants (Figure 7C).

**Figure 7:**
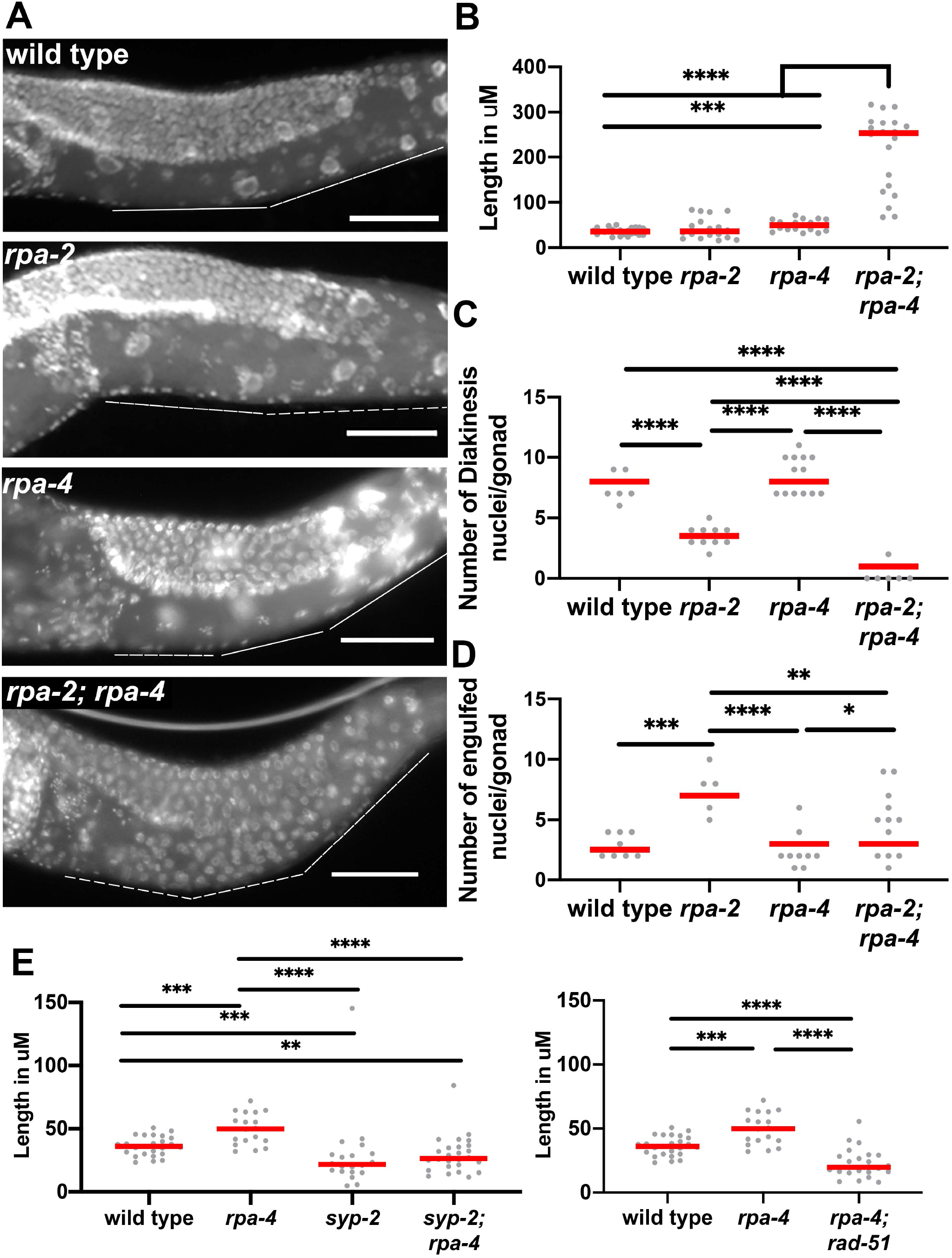
Double deletion of *rpa-2* and *rpa-4* leads to meiotic progression defects, with defects in apoptosis. A) Representative images of gonads in Carnoy’s fixed whole worms. (scale bar= 50μm) B) Number of CED-1::GFP engulfed nuclei in 3-day-old worms. C) Number of diakinesis oocytes in each mutant background. D) Number of CED-1::GFP engulfed nuclei in IR treated and mock-treated worms 24 hours after treatment. E) Length of the gonad from the bend to the first diakinesis nucleus in microns. (Mann-Whitney tests performed, where p-values are represented as ****=<0.0001, ***=<0.001, **=<0.01, and * =<0.05)

In the germline, physiologic apoptosis is used as a mechanism for clearing half of the meiotic nuclei so their metabolite content can be supplied to the few oocytes targeted for fertilization (31). Apoptosis is also used for removing damaged cells, but this is not thought to contribute to apoptosis under normal growth conditions. The accumulation of pachytene nuclei in *rpa-2; rpa-4* double mutants can be attributed to defects in apoptosis. To test this hypothesis, we examined the level of apoptosis in the three mutants using CED-1:GFP reporter. CED-1 is a transmembrane receptor of the *C. elegans* germline that mediates engulfment of apoptotic nuclei (35). While *rpa-4* had no effect on apoptosis, *rpa-2* mutants exhibited a notable increase in apoptosis, regardless of whether the total number of apoptotic nuclei was normalized or not to the number of nuclei (Figure 7D and S5). This increase is expected since in the absence of RPA-2, HR is abrogated and nuclei with unrepaired DSBs accumulate. Consistent with RPA-4 playing a role in promoting apoptosis, *rpa-2; rpa-4* double mutants displayed wild-type levels of engulfed nuclei. These data indicate that meiotic progression defects in *rpa-2; rpa-4* double mutants are attributed to the loss of the apoptotic response.

In light of the opposing function RPA-4 plays in RAD-51 and RPA-1 focus assembly, the genetic interaction between *rpa-2* and *rpa-4* may be interpreted as a requirement of RPA-4 for apoptosis in the presence of extensive DNA damage found in *rpa-2* mutants. If true, removal or *rpa-4* in a different genetic background that increases DNA damage will phenocopy the *rpa-2; rpa-4* double mutant phenotype. SYP-2 is part of the synaptonemal complex which pairs homologous chromosomes in meiosis I, and mutants of *syp-2* have increased apoptosis due to delay in HR and abrogation of inter-homolog DSB repair. In *syp-2; rpa-4* mutants, extension of pachytene was not observed. DSB repair defects in *syp-2* mutants are different than those of *rpa-2* mutants, since the former contains ssDNA bound by RAD-51 capable of stand invasion, while the latter does not. To further test our hypothesis, we combined *rpa-4* deletion with *rad-51* mutants that contain resected ssDNA and no functional HR. However, *rpa-4; rad-51* double mutants were indistinguishable from *rpa-4; syp-2* double mutants, or *rpa-4* single mutants (Figure 7E). Taken together, these data indicate that under normal growth conditions, RPA-4 is responsible for apoptotic signal that is are upstream of RAD-51 binding.

### RPA-4 acts to promote germline apoptosis in aging germlines and following DNA damage in a CED-3 independent pathway

RPA-4 is not required for apoptosis under normal growth conditions in young adult worms (Figure 7D). However, RPA-4 is found in low levels under these conditions and is only recruited to DNA damage following exogenous DNA damage. It is therefore possible that RPA-4 promotes apoptosis in challenging conditions. Apoptosis has been shown to increase in aging worms (51). Therefore, we examined the localization of *rpa-4* foci in 3-day-old worms and found a significant increase in RPA-4 foci numbers (Figure 8A). Unlike what is found in young adults, RPA-4 foci numbers were partially dependent on SPO-11 induced breaks as they were almost abolished in the *spo-11* mutants. If the elevation of RPA-4 foci in aged worms was important for apoptosis, then apoptosis in these worms will be *rpa-4* dependent. Indeed, apoptosis levels in 3-day-old adults were reduced in *rpa-4* mutants compared to wild type(Figure 8B). An effect on oocyte numbers was also observed following exposure to ionizing radiation and was mildly suppressed in *rpa-4* mutants (Figure 8C). When engulfed nuclei of wild-type germline were examined for colocalization with RPA-4, they showed no increased preference for RPA-4 (Figure 8D). These data altogether indicate that RPA-4 plays a role in apoptosis under challenging conditions.

**Figure 8:**
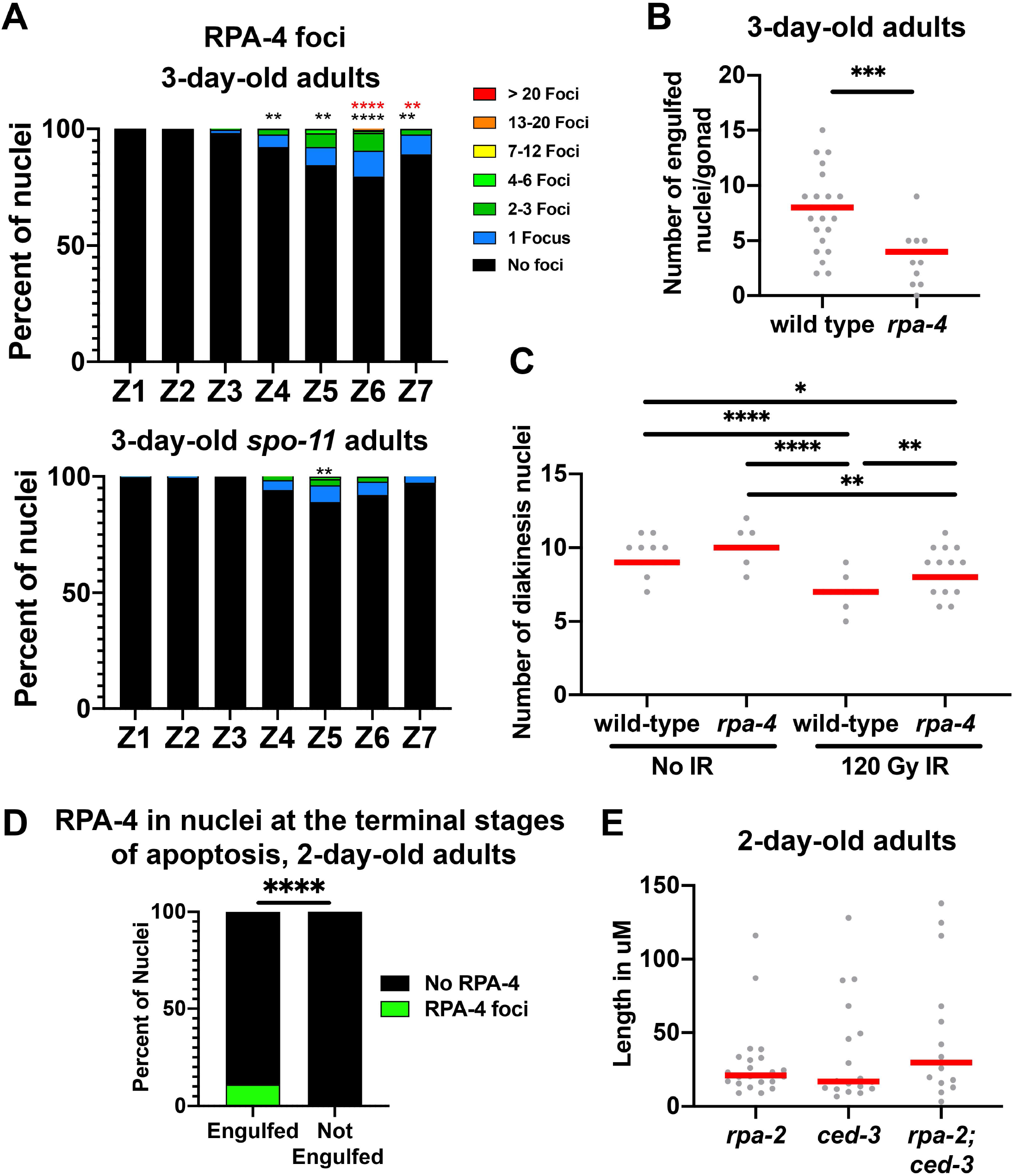
RPA-4 foci appear in greater abundance in aging worms, effecting apoptosis signaling through a CED-3 independent pathway. A) Percent of nuclei with indicated amount of FLAG::RPA-4 foci. Black asterisks are comparison with wild type 1-day-old adult worms. B) Number of CED-1::GFP engulfed nuclei in wild type and *rpa-4* mutant 3-day-old worms. C) Number of diakinesis oocytes in wild type and rpa-4 worms in 24 hours post-IR and mock-IR. D) Presence of FLAG::RPA-4 or FLAG::RPA-2 positive nuclei in or outside of CED-1::GFP engulfment (red-comparison to *spo-11* 3 day old, black, comparison to wild type 1 day old). E) Length of gonad with pachytene-like nuclei after the bend in indicated mutant backgrounds in 2-day-old adults. (Mann-Whitney tests performed, where p-values are represented as ****=<0.0001, ***=<0.001, **=<0.01, and * =<0.05)

Next we aimed to identify where RPA-4 acts in the apoptotic pathway. Mitogen-activated protein kinase (MAPK) has been identified as an important factor influencing germline apoptosis, and mutants of *mpk-1* show germline arrest prior to diplotene (37, 52). We stained for activated MAPK (di-phosphorylated MPK-1) and found that *rpa-2; rpa-4* double mutant gonads displayed normal MAPK staining (Figure S6). This indicates that “licensing” for apoptosis is occurring in *rpa-2; rpa-4* double mutants, but commitment to apoptosis is altered. These data suggest that RPA acts downstream of MAPK.

If RPA-4 acts though the canonical apoptosis pathway, then removal of apoptosis in the *rpa-2* mutants should recapitulate the extended pachytene phenotype of *rpa-2; rpa-4* double mutants. CED-3 is involved in initiating apoptosis in the germline for physiological apoptosis and in response to DNA damage (53, 54). The *rpa-2; ced-3* double mutants had germline progression that was indistinguishable from *rpa-2* or *ced-3* (Figure 8E). It is known that CED-3-independent apoptosis occurs in the gonad, and these data indicate that RPA-4 could interact with one of these apoptotic mechanisms (55). Taken together, RPA-4 is involved in apoptosis, through yet unidentified CED-3-independent mechanism.

## Discussion

These data provide insight into the roles of alternative RPA complexes (RPA-1/RPA-2 and RPA-1/RPA-4) in *C. elegans* meiosis. We demonstrate that RPA is essential for replication and recombination as is seen in other organisms, and we identify an additional role of RPA in regulating germline development. This work offers the first look at the role of RPA-2 and RPA-4 in the *C. elegans* germline, as well as a comprehensive investigation of RPA-1. Our data suggests that RPA-1 and RPA-2 are essential for normal germline replication, and that RPA-4 acts as part of a quality control mechanisms promoting germline apoptosis. Our works suggests that RPA-4 evolved to provide a response for conditions in which DSB repair is impaired (older worms) or challenged by excessive damage (ionizing radiation/HU). Despite the huge similarity between RPA-4 and RPA-2 these proteins evolved to play different and antagonistic functions in DSB repair.

### RPA-1 and RPA-2, not RPA-4, are critical for normal germline replication and HR repair

Previous reports have demonstrated that the large, medium, and small RPA subunit are essential for replication and recombination across organisms (39). We examined the function of the known RPA subunits in *C. elegans*: one RPA1 and two RPA2-like subunits. Our biochemical studies suggested that no trinary complex is formed between these known subunits. This does not exclude the presence of a third, yet unidentified subunit that performs a function similar to RPA3. This is formally possible as canonical metazoan RPA is a trimeric complex (56). Instead all pairwise combinations are possible. While RPA-1/2 and RPA-1/4 may be expected, based on studies in other organisms, the RPA-2/4 dimer is unexpected and may explain the genetic interactions between RPA-2 and RPA-4 (see below).

Our data is consistent with a role for *C. elegans* RPA-1 and RPA-2 in replication (likely as RPA-1/2 complex). However, while RPA-1 is essential for replication, RPA-2 is not. This suggest that RPA-1 can bind ssDNA and facilitate replication without the need to bind RPA-2. Instead, RPA-1 activity is enhanced by RPA-2 binding. Our localization data is consistent with this model, as RPA-1 can localize to mitotic germline nuclei in the absence of *rpa-2*, albeit in lower levels than wild type. This observation is perplexing, as it was assumed that in metazoans RPA is an obligatory trimeric complex. In other organisms it was shown that RPA3 is recruited to the RPA complex by physically interacting with RPA1 and RPA2. The ability of RPA-1 to promote replication by itself may also explain why a putative RPA-3 has not yet been described or is possibly missing. Despite the sequence similarities, we have seen no evidence for RPA-4, the RPA-2 ortholog, playing a role in replication. *rpa-4* mutants show no phenotypes related to replication defect and RPA-4 does not extensively localize to mitotic nuclei even under HU stress.

The second conserved function of the RPA complex is in promoting HR repair by binding to ssDNA following resection, which is essential for RAD-51 recruitment. Our data is consistent with RPA-1 and RPA-2 playing a conserved role in this process. Unlike what is found for replication, RPA-2 plays an essential role in HR. This may be a result of an interaction of RPA-2 with resection factors, or with proteins that allow for loading of RAD-51 (24, 48). In its absence HR fails and despite the ability of RPA-1 to localize to the ssDNA formed, it cannot support efficient RAD-51 recruitment. In *rpa-2* mutants the DSBs formed in meiosis are repaired by cNHEJ and TMEJ while HR is abrogated. This data suggest that RPA-1-ssDNA filament is more permissive to replication then to HR. As with replication, RPA-4 doesn’t seem to play a role in HR under normal conditions.

### RPA-4 acts in promoting apoptosis in challenging conditions

Under normal growth conditions (young adults with no exogenously induced damage), deletion of *rpa-4* does not affect any phenotype indictive of a function in DNA damage repair. The localization pattern of RPA-4 in these conditions indicates that very few nuclei require RPA-4’s function and that RPA-4’s foci are not localized to SPO-11 induced breaks. However, under challenging conditions (older worms/γIR/HU), RPA-4 is found in more nuclei and is promoting apoptosis. We interpret these data as indicating a role of RPA-4 in targeting a subset of nuclei, ones with unrepairable DSBs, to apoptosis.

Our data suggested that RPA-4 acts in a non-canonical apoptosis pathway that is CED-3 independent. RPA-4 is also placed downstream of MAPK signaling. The fact that RPA-4 foci are found throughout pachytene, and not just in late pachytene argues that RPA-4 acts prior to the implementation of apoptosis (which is restricted to late pachytene). RPA-4 is recruited to DNA damage foci following RPA-1. We propose that following DNA damage RPA-1/2 localizes to DSBs. While some DSBs can be promptly repaired, others may be more challenging to repair. In normal conditions these may include DSBs generated by replication errors but in aging worms altered DNA repair processes may also channel SPO-11 induced DSBs to this pathway. The identified nuclei are then marked by RPA-4, and these nuclei move to late pachytene, where MAPK signaling occurs and apoptosis is executed.

### RPA-4 and RPA-2 function antagonistically in regulating RPA-1 focus formation

RPA-2 performs a canonical function in DSB repair by promoting RPA-1 activity. In its absence, RPA-1 focus formation is impaired, but the effect is much larger on RAD-51 focus formation, indicating that in addition to facilitating RPA-1 engagement with DNA, RPA-1/2 complex can support RAD-51 focus formation more efficiently than RPA-1 by itself. RPA-2 not only promotes RPA-1 function but also inhibits RPA-4 focus formation. This may be an indirect action due to RPA-1/2 being a complex that more easily associates with ssDNA than RPA-1, outcompeting for RPA-4 binding. Alternatively, RPA-2 can effect RPA-4’s ability to bind ssDNA. Indeed, in our single molecule experiments we have shown that RPA-1 binds RPA-4 forming a stable complex. This interaction may negatively and directly regulate RPA-4 binding to ssDNA.

Despite high similarity between RPA-1 and RPA-4, RPA-4 not only plays a completely different function in HR, but also an antagonistic role to RPA-2. Instead of promoting RAD-51 focus assembly, RPA-4 attenuates it. Our MIR studies show that RPA-4 is recruited to DSBs following RPA-1. We propose that RPA-4 recruitment to RPA-1 foci can disassociate RPA-1 to an extent that ssDNA doesn’t contain enough RPA-1 to support RAD-51 loading, or RPA-4 prevents the displacement of RPA-1 by RAD-51. As a result, the nucleus is committed to apoptosis. Since RPA-4 is recruitment is delayed, this can provide enough “buffer time” for the nucleus to load RAD-51 and repair the DSB through HR. On challenging DNA damage repair is delayed, which results in RPA-4 recruitment and apoptosis.

## Methods

### Strains: and maintenance

Worms were maintained on NGM plates with lawns of OP50 *E. coli* at 20°C. Strains used for experiments include N2 (Bristol), and contained the following alleles in the N2 genetic background:

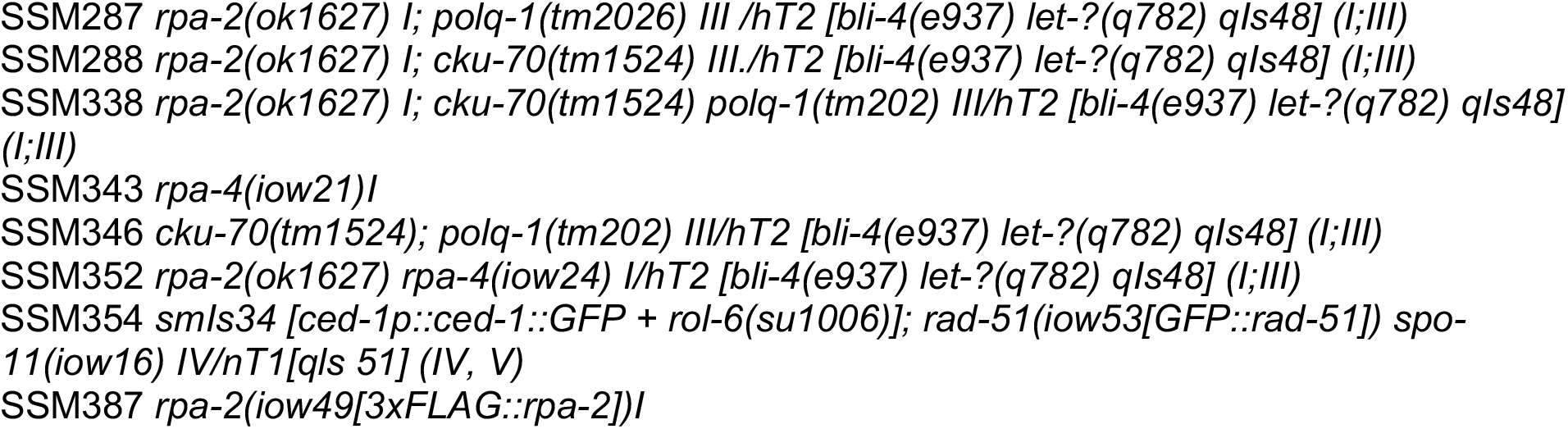
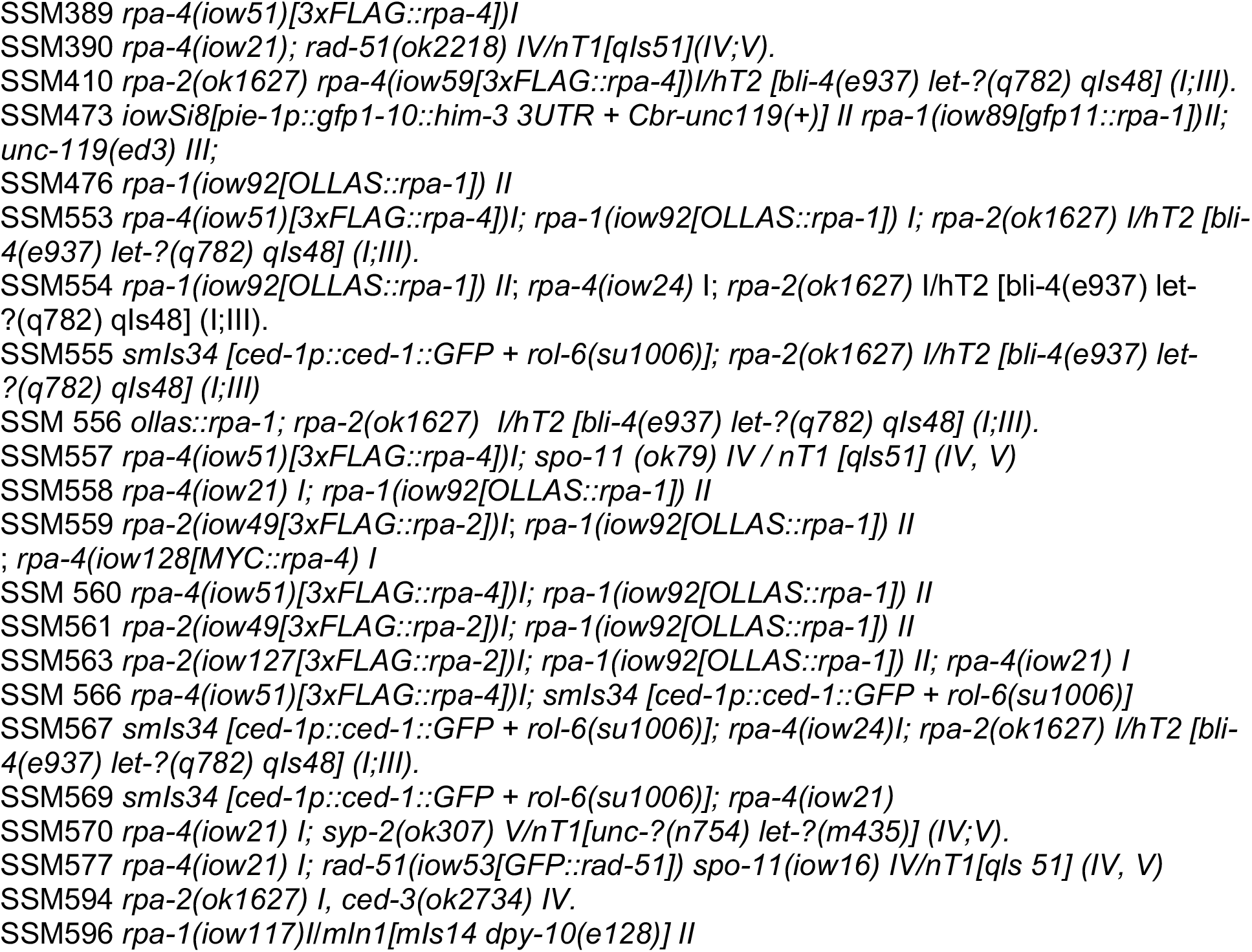

### Viability

L4 worms were singled onto NGM plates containing a small (1 cm^3^) lawn of OP50 *E. coli*. P0s were transferred to new plates twice a day for 4 days. Immediately after transfer, eggs were counted, and adults were counted 4 days later.

### CRISPR

CRISPR/Cas9 was used to create the following strains for this publication:

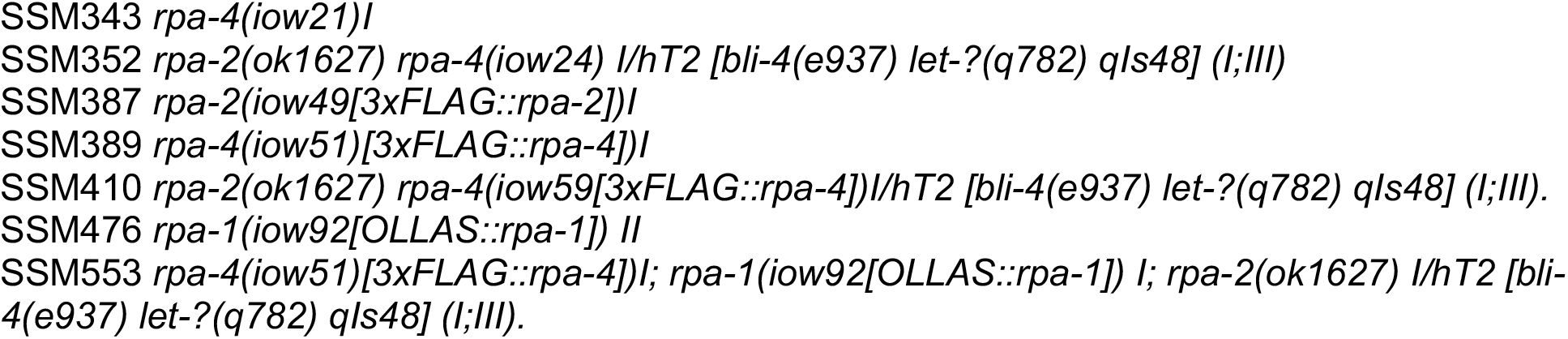
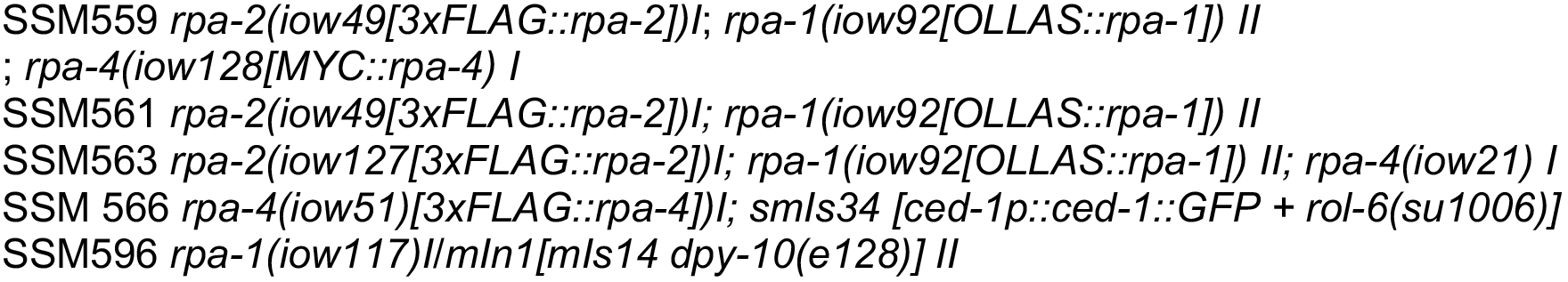

Micro-injection of 1-day-old adult worms was performed on 3% agarose pads, afterwards collected on a single NGM plate, and isolated to individual OP-50 seeded plates the following morning. Plates were screened for the *rol* or *dpy* phenotypes generated by *dpy-10* point mutation introduced by co-CRISPR marker, adopted from (57). Wild-type F1s were isolated to individual plates for insertion screening by PCR and sequencing. tracrRNA, and crRNAs were obtained from IDT and mixed in the following concentrations: 14.35 μM Cas9-NLS (Berkeley MacroLab), 17.6 μM tracrRNA (IDT), 1.5 μM *dpy10* crRNA (IDT), 5 μM *dpy10* ssODN (IDT), 16.2 μM of target crRNA (IDT), and 6 μM of target ssODN (IDT)). ssODNs and crRNA used include the following:

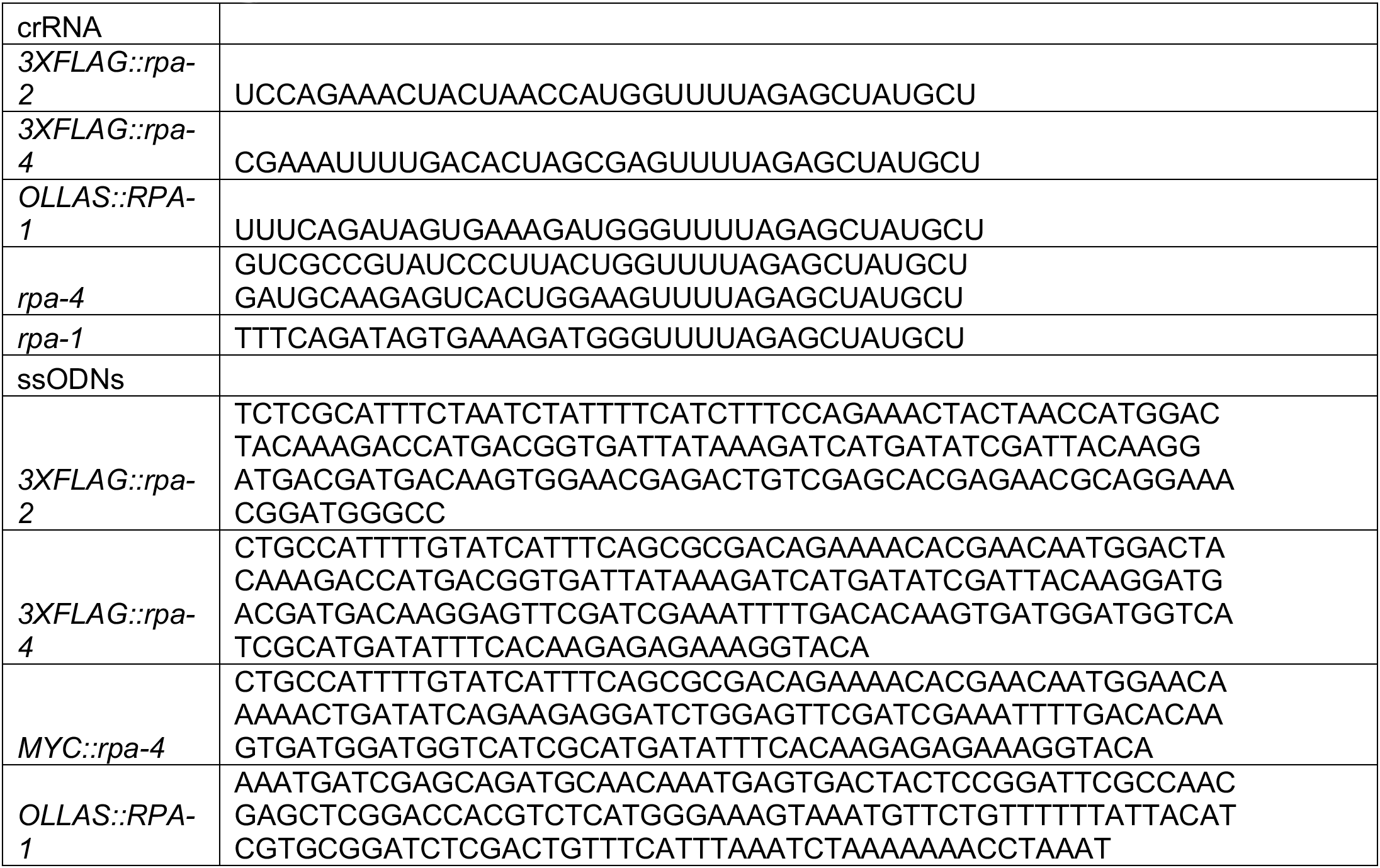

### Western

Protein samples had SDS urea lysis buffer and 2-mercaptoethanol added, and were immediately boiled for 5 minutes. After boiling, samples were vortexed for 2 minutes before being transferred to ice. Samples were run on a 10% SDS Express plus PAGE gel (#M01012; GenScript) using SDS-MOPS buffer. Protein was transferred to a nitrocellulose membrane in tris-gylcine buffer, washed briefly in 1XPBS-tween, and blocked with 5% dehydrated milk in 1XPBS for 1-2 hours. was used as wash buffer. Primary antibody was diluted in 1XPBS-tween+milk overnight at 4C. The following morning, the membrane was washed 3 times using 1XPBS-tween before 2 hour incubation with secondary antibody, followed by three more washes. Using WesternBright ECL (#K-12045-D20; Advansta), blots were exposed using the LI-COR Odyssey Infrared Imaging System. Antibodies used were as follows: Rat α-ollas (NOVUS, NBP1-06713, 1:2000), Peroxidase AffiniPure Goat α-Rat IgG (H+L) (1:5000)

Mouse anti-flag 1:2000 (Sigma F1803), Goat α-Mouse IgG (Kappa light chain) HRP (1:1000)

### Antibody staining and image acquisition

10-20 worms were dissected using a #10 razor blade of in M9 on a coverslip. Immediately after gonads were extruded, coverslip was transferred to a positively charged slide and flash frozen on dry ice. Preparation of *ced-1::gfp* worms was performed such that slides were kept in the dark for as long as possible. The coverslips were removed and slides were in methanol for 1 minute, and a 30 minute fix (25 minutes for *ced-1::gfp* worms) in 4% paraformaldehyde (Alpha Aeser) made from 37% stock. Slides were dipped in 1XPBST for 10 minutes, and then incubated with 0.5%BSA in 1XPBST for 1-2 hours. Afterwards, they were incubated with primary antibody over-night at room temperature. The following day slides were washed in 1XPBST three times, incubated with secondary antibody for 2 hours at room temperature in the dark, and then washed in 1XPBST. Slides were then incubated in the dark with a 4′,6-diamidino-2-phenylindole (DAPI, 1:10,000 of 5mg/ml stock in 1XPBST), followed by a final wash in 1XPBST. Slides were sealed with VECTASHIELD (Vector Laboratories) and stored at 4°C. Antibodies were used in the following concentrations:

Mouse α-flag (Sigma F1803, 1:500), Alexa Fluor 488 α-mouse (1:500)
Mouse α-flag (Sigma F1803, 1:500), DyLight 550 α-mouse (1:500)
Rabbit α-rad-51 (1:30000), Alexa Fluor 488 α-mouse (1:500)
Rat α-ollas (NOVUS NBP1-06713, 1:75), Alexa Fluor 594 α-rat (1:2000)
Rabbit α-ollas (Genscript A01658,1:1000), Alexa Fluor 488 α-rabbit (1:500)
Rabbit α-PCN-1 (Gift from M. Michael,1:13000), Alexa Fluor 488 α-rabbit (1:500)
Rabbit α-SUN-1 (Novus, 1:1000), Alexa Fluor 488 α-rabbit (1:500)
rabbit α-PH3 (1:5000), Alexa Fluor 488 α-rabbit (1:500)
Mouse Anti-MAPK-YT (Sigma) 1:400, Alexa Fluor 488 α-mouse (1:500)

All images were taken using the DeltaVision wide-field fluorescence microscope (GE lifesciences) with 100×/1.4 NA oil Olympus objective. Images were deconvolved with softWoRx software (Applied Precision) unless otherwise noted.

### Carnoy’s and ethanol fix

For whole-worm mutant comparisons, and CED-1::GFP engulfment analysis, worms of the indicated ages were placed on an uncharged slide (Surgipath Leica) inside a drop of M9. The M9 was removed using Whatman filter paper, and 8 microliters of 95% ethanol (Millipore) (for CED-1::GFP analysis) or Carnoy’s fix(60% ethanol, 30% chloroform, 10% glacial acetic acid))(for gonad length comparison) was added to the worms, and then allowed to evaporate. To preserve the worms and stain chromatin, 9μl of Vectashield with DAPI was added to the slide, and a #1.5 coverslip was placed on top, before sealing with acrylic nail hardener

### Whole worm imaging

Images were taken on a Leica DMRBE microscope using a 10X/0.30 PL FLUOTAR objective. A QIClick (QIMAGING) camera captured images using Q-Capture software. Scale bars in the whole worm images represent 50μm.

### Nuclear volume

16-bit non-deconvolved images of SUN-1 antibody stained gonads were taken with the DeltaVision microscope at 100X magnification in 0.1 micron slices. The area at the largest part of each nucleus(a), and the number of slices(b) from top to bottom of each nucleus was measured for 5 nuclei and from at least 3 gonads for each genotype. Using the following formula, volume was estimated based on an ellipsoid volume calculation:
*V*=4/3*π(a*/*π)(b* × .5)

### Immunoprecipitation

Worms were chunked to 10 OP50 seeded NGM plates, and allowed to grow for 3-4 days before being washed off with M9 into a 15mL conical tube. Worms were pelleted and washed with M9 until bacteria was removed (~5 times). Lysis was performed in a Precelly® machine with equal volumes of Pierce® IP Lysis buffer(Thermo, #8778). The resulting slurry was pelleted, and the supernatant was added to a tube containing 50 microliters of Anti-FLAG^®^ M2 Magnetic Beads (Sigma, #m8823). This mix was allowed to incubate on a rotating mixer at 4C overnight. After overnight incubation, the beads were pelleted using a magnetic stand, and the supernatant was collected. The beads were washed 5 times using PBS+ ROCHE cOmplete™ Protease Inhibitor Cocktail tablet. Bound proteins were eluted with Thermo Gentle Ag/Ab elution buffer (#21027) and Thermo IgG Elution Buffer (#21004) for 5 minutes each.

### Quantification of engulfment using CED-1::GFP

Images of *ced-1::gfp* were analyzed using softWoRx software (Applied Precision), and engulfed nuclei were scored only if there was complete engulfment.

### HU treatment

1-day-old worms were transferred to OP50 seeded and sterilized 40mM HU plates that were made the same day. Worms were allowed to incubate at 20C for 20 hours before transfer to OP50 seeded plates. To determine viability, worms were counted for 3 days as described before.

### Gamma IR

L4’s were transferred to OP50 seeded NGM plates, then exposed to a cesium source 24 hours after transfer. For generating DSBs and analysis of RPA-4 focus formation, 120 gray was used, and for viability of *rpa-4* mutants, 100 gray was used. For RPA-4 localization, gonads were dissected 6 hours after exposure to IR. For gonad length and apoptosis analysis, worms were allowed to recover for 24 hours after treatment.

### Length measurements

Using FIJI, length intensity measures were gathered by drawing a line through the middle of the gonad from either the bend of the gonad to the first diakinesis nucleus, from the distal tip to the last diakinesis nucleus, depending on the analysis.

### Microirradiation

Microirradiation was performed and analyzed according to the protocol outlined in Koury et al., 2018 with the following alterations: 1) 2 worms were placed on each live imaging slide, 2) for data presented in Figure S4, imaging was performed at 2 minute intervals for 1 hour, and 3) for data presented in Figure 6B-C, imaging was performed at 2 minute intervals for 45 minutes with 10% light source intensity, and 250ms exposure in the GFP channel. All experiments were performed at 15% pulse intensity. For gfp11::rpa-1; gfp1-10, worms were grown at 25°C, L4s were selected from these plates, and then allowed to grow at 20°C until use in the experiment the next day as 1 day old adults.

### Antibody labeling for single-molecule analysis

The following antibodies and reagents were used:

Mouse α-flag (Sigma F1803);
Rat α-ollas (Novus Nbp1-06713);
Rabbit α-Myc (Sigma C3956);
EZ-Link NHS-Biotin (ThermoFisher Scientific 20217);
Cy3 Mono-Reactive NHS Ester (Millipore Sigma PA13105);
Cy5 Mono-Reactive NHS Ester (Millipore Sigma PA15101);

An Amersham protocol for conjugating NHS reactive reagents was used but modified with the following steps. Zeba desalting columns (0.5ml) were spun at 1500g to remove storage buffer. The columns were equilibrated with 1M NaHCO_3_. After the columns were equilibrated 100uL of antibody solution was added. The antibodies were spun at 1500g for 2 minutes. The solution was recovered, and protein concentration was measured using Nanodrop-2000. The equation to calculate the protein concentration at 280nm is A=εcL, where A is the absorbance at 280nm, ∊ is the molar extinction coefficient, for the antibodies used this was approximately 170,000M^−1^cm^−1^, and L is the path length which is 1 cm. The volume of NHS reagent (Cy3, Cy5 or NHS-Biotin) was calculated so that there would be 20:1 reagent/antibody ratio in the reaction mix. The reagent and antibody were then combined in a 1.5ml centrifuge tube, mixed and allowed to incubate at room temperature for 1 hour. After incubation another spin column was prepped using the same method as above, however the spin columns were equilibrated with DPBS. The antibody/dye solution was spun down and the flow through was collected. This solution was then measured for protein concentration and respective dye-to-antibody ratio.

### Preparation of *C. elegans* lysate for single-molecule studies

Worms were grown on OP50 lawns and chunked 4 days before being rinsed from plates using M9. Pierce™ IP Lysis Buffer (Thermo) was prepared with 1 tab of protease inhibitor for 10mL of buffer. After rinsing worms from plates using M9, worms were pelleted by centrifugation at 1600×G for 20 seconds, and supernatant was removed. Lysis buffer was added 1:1 to worm pellet. One mL of this mix was placed into Precellys lysis tubes (VK05). Lysis was performed using a Precellys machine for 15 seconds, before putting the sample on ice. This procedure was repeated 3 more times, and samples were spun at 2000xG for 5 minutes at 4°C. Supernatant was collected and stored at 4°C.

### SiMPull

Reagent Buffer (RB) was used for diluting non-fluorescently labeled antibodies or cell lysate. The RB consists of (1mg/ml BSA, 0.01% v/v Triton x-100, and DPBS (Gibco)). Imaging Buffer (IB) was used to dilute fluorescently labelled antibodies and for washing flow cells when fluorescent reagents were present. IB consists of (25% RB v/v, Gloxy (0.04mg/ml), 0.8% glucose and Trolox). Trolox was made by using 12 mM Trolox (6Hydroxy-2,5,7,8-tetramethylchromane-2-carboxylic acid, Sigma-Aldrich, MO, USA; 238813–1G) dissolved in 12 mM NaOH. The solution was then rotated under a fluorescent light (Sylvaia FM13W/835) for 3 days or until the absorbance at 400 nm is approximately 0.119. Gloxy was made by mixing (40mg/ml) catalase in T50. Then 10ul of this solution is mixed with 90ul (10mg/ml) glucose oxidase. Buffers were stored in 4°C. Buffers were allowed to reach room temperature prior to using in experiments.

SiMPull experiments (47) were carried out using homebuilt total internal fluorescence microscopy system (TIRFM). The system and the TIRFM flow cells are described in (58). A diode-pumped solid state (DPSS) green laser (532 nm, Coherent, CA, USA) was used to excite the Cy3 dye. A diode-pumped solid state (DPSS) red laser (645 nm Coherent, CA, USA). The laser power output was set to 45 mW for all images. A dual band pass filter (Semrock, NY, USA; FF01-577/690) was used to filter scattered light in the optical path. The fluorescence was collected using a Chroma ET605/70m filter. Movies were taken using an electron-multiplying charge-coupled device (EMCCD) camera (Andor, MA, USA; DU-897-E-CSO-#BV). Exposure setting was 100ms for all movies. During recording background was set to 400, and correction to 1200. The gain was set to 295 for all movies.

*C. elegans* lysates from the wild type animals and animals expressing tagged versions of RPA genes were stored in 4° or kept on ice during experiment until used in the flow cell. After flow cell assembly and TIR acquisition, RB was used to wash the chamber. Neutravidin (0.5mg/ml) was flowed through the chamber and allowed to incubate for 3-5 minutes. After incubation the chamber was washed again with RB. Biotinylated antibody was then flowed into the chamber. For all instances, the biotinylated antibody was diluted to approximately 10^−9^ M. The biotinylated antibody was allowed to incubate in the chamber for 10 minutes. After incubation the excess antibody was removed by flowing RB through the chamber. Cell lysate from stock was flowed through the chamber and was allowed incubate for 30 minutes. Next Cy3 and Cy5 labelled antibodies were mixed in IB + 0.8% glucose to a concentration of approximately 10^−11^ M. IB labelled-antibody solution was then flowed through the chamber, the lasers are turned off to ensure that during incubation there was minimal photobleaching of the fluorophores. The labelled antibodies were allowed to incubate with the lysate for 30 minutes. The chamber is then washed with 200 μl of IB + 0.8% glucose. Several videos are then taken in repetition in unique fields of view (FOV). A second wash of IB + 0.8% glucose is flowed through the chamber and subsequent videos in unique (FOV).

Data were quantified using an ImageJ-FIJI plugin (TrackMate). The settings that were used to extract the values were as follows. The difference of gaussian (DoG) segmenter was used to identify particles. Approximate particle diameter was set to 0.04 inch, and threshold was set to 2. No initial thresholding was used. No filters were set for spots. The Linear Assignment Problem (LAP) Tracker was used to track particles across each image. The maximal distance for frame to frame linking was set to 0.01 inches and a maximum track segment gap closing value of 0.01 inch and 2 frames was used. The results were exported, and the data was visualized using Prism 8.0.

### Statistical analysis

All data was analyzed with Graphpad Prism 7 software. When multiple groups were compared, and non-parametric data used, Kruskal-Wallis tests were applied, with multiple comparisons using the two-stage linear step-up procedure of Benjamini, Krieger and Yekutieli to control for false positives. When q-values were significant (q<0.05), statistical significance was reported from Mann-Whitney analysis. For comparison nuclear volume in the PMT, data was normally distributed (according to Anderson-Darling, D’Agostino & Pearson, Shapiro-Wilk, and Kolmogorov-Smirnov tests), but variance was not equivalent, therefore a T-test with Welch’s correction was applied reported (Figure 1H). For complete gonad length measurement, data was normally distributed with equivalent variances, therefore T-test was applied and values reported (Figure S2B). For SiMPull data comparison 2-way ANOVA in Prism 8.0 was used.

## Supporting information

Hefel Sup data

## Data availability

Strains are available upon request. The authors state that all data necessary for confirming the conclusions presented in the article are represented fully within the article.

## Acknowledgments

Some strains and clones were kindly provided by the *Caenorhabditis* Genetics Center, which is funded by the National Institutes of Health (NIH) Office of Research Infrastructure Programs (P40OD-010440), and the *C. elegans* Reverse Genetics Core Facility at the University of British Columbia, which is part of the International *C. elegans* Gene Knockout Consortium. We thank the National Bioresource Project for the Experimental Animal “Nematode C. elegans”, Japan for providing alleles for this study. We thank M. Michael for the PCN-1 antibody and E. Koury for her help with experiments. We thank M. Wold for helpful discussions and reading the manuscript. We acknowledge the staff at the University of Iowa College of Medicine and Holden Comprehensive Cancer Center Radiation and Free Radical Research (RFRR) Core for radiation services. The RFRR core facility is supported by funding from NIH P30 CA086862. This work was funded by NIH grant number 1R01 GM-112657 (to S.S.) and R35GM131704 (to M.S.).

