## Supplementary material for "RPA complexes in *Caenorhabditis elegans* meiosis; unique roles in replication, meiotic recombination and apoptosis": Hefel Sup data

Figure S1

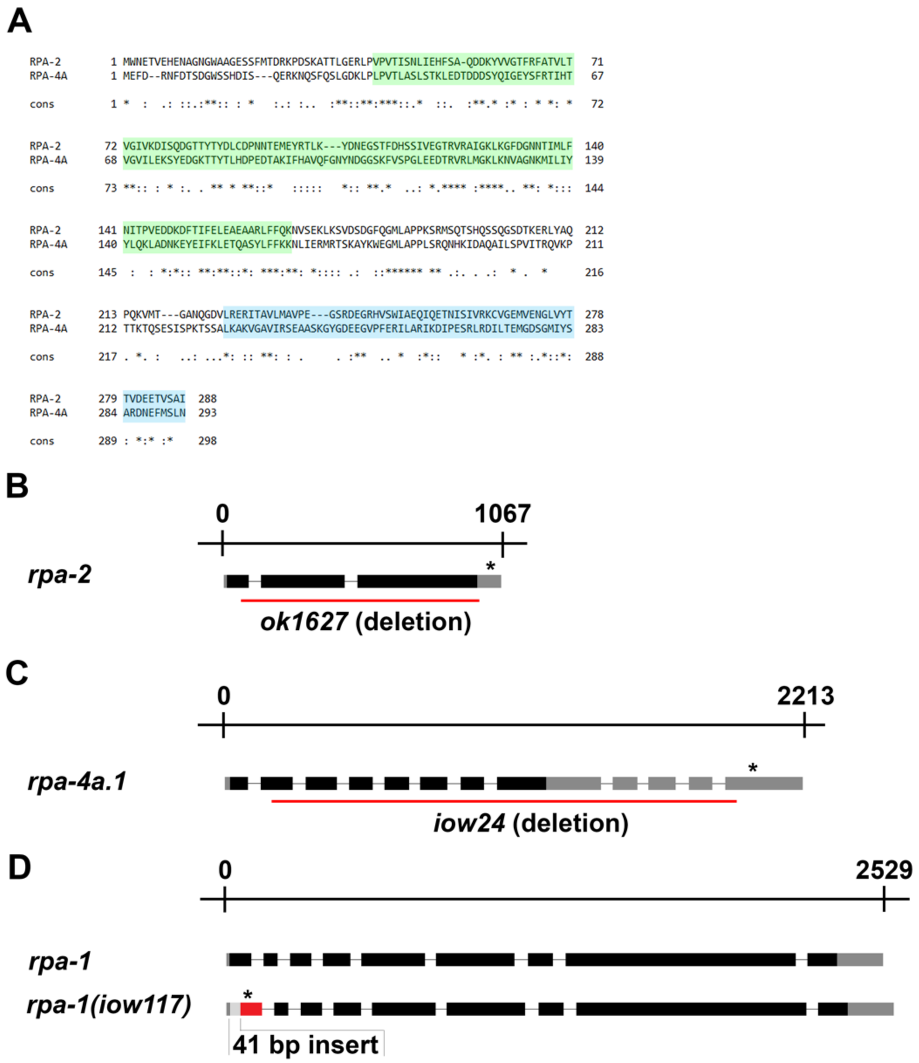


**Figure S1: Alignment of RPA-2 and RPA-4 protein sequences, and mutant illustrations.** A) ClustalW alignment of RPA-2 with RPA-4. Green box indicates OB-fold domain. Blue box indicates winged-helix domain. B-D) Illustrations of the corresponding genes and mutations used in this manuscript. Black lines represent introns. Black boxes represent exons. Grey boxes represent untranslated regions. Asterisk represents premature stop codon.

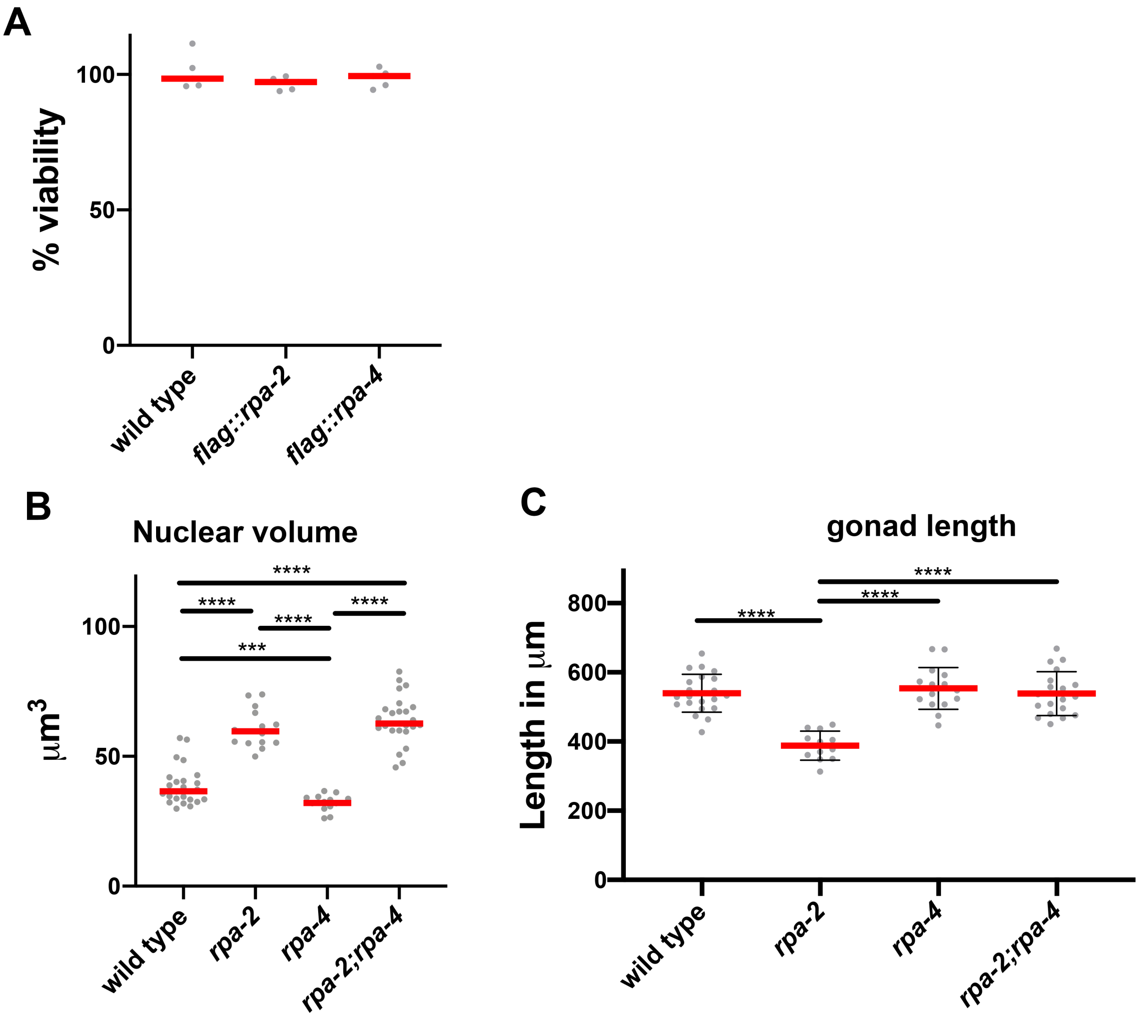


**Figure S2: Viability of tagged proteins and mutant phenotypes of germline size.**
A) Percent viability of eggs laid in wild type and FLAG tagged *flag::rpa-2* and *flag::rpa-4* lines. B) Nuclear volume of mid-pachytene nuclei. C) Length of gonad from the distal tip to the end of diakinesis. (Mann-Whitney tests performed, where p-values are represented as ****=<0.0001, ***=<0.001, **=<0.01, and * =<0.05)

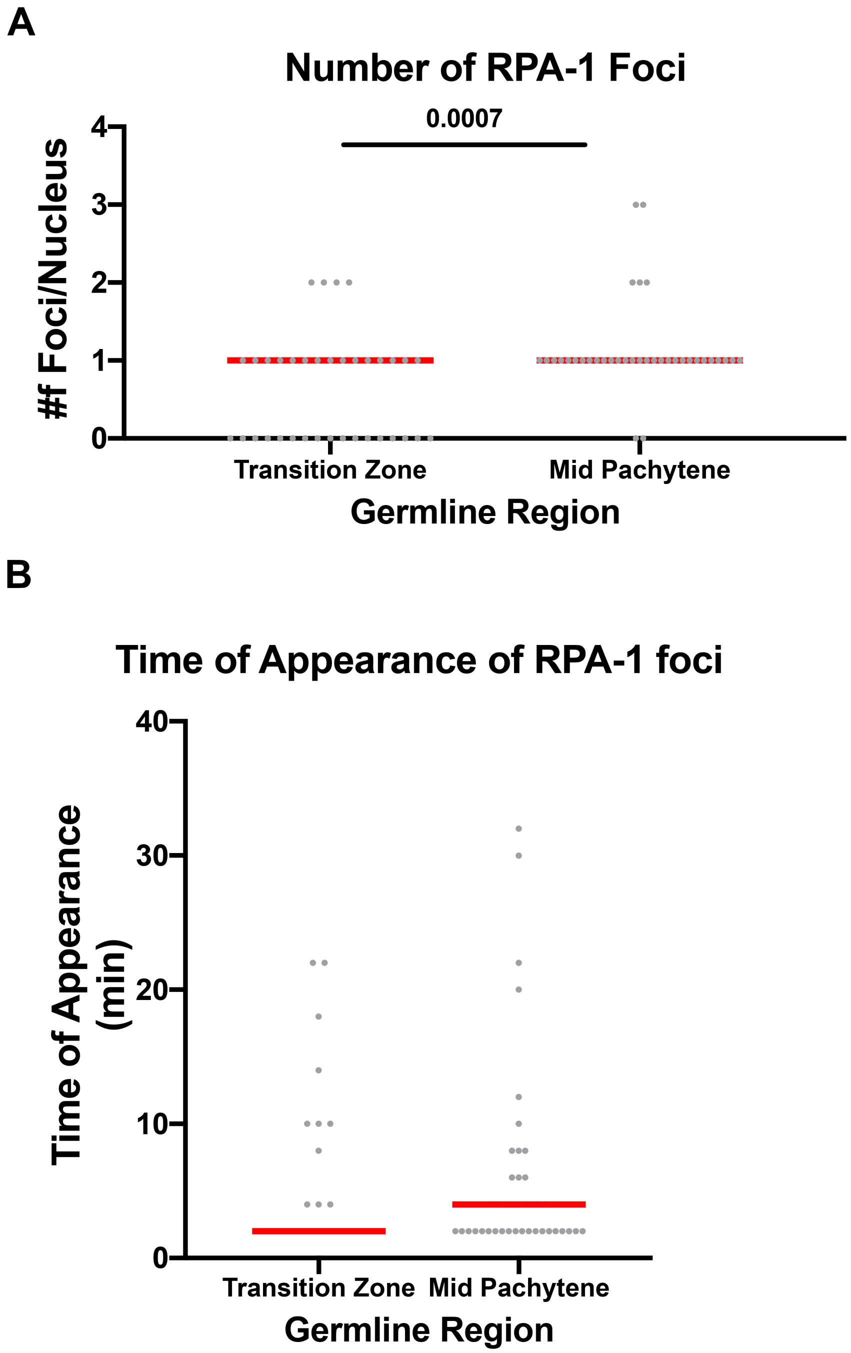


**Figure S3: GFP11::RPA-1 focus appearance following UV laser micro-irradiation.**
A) Number of GFP11::RPA-1 foci that appear in the indicated regions in each nucleus. B) Time of appearance for each focus. (Mann-Whitney tests performed)

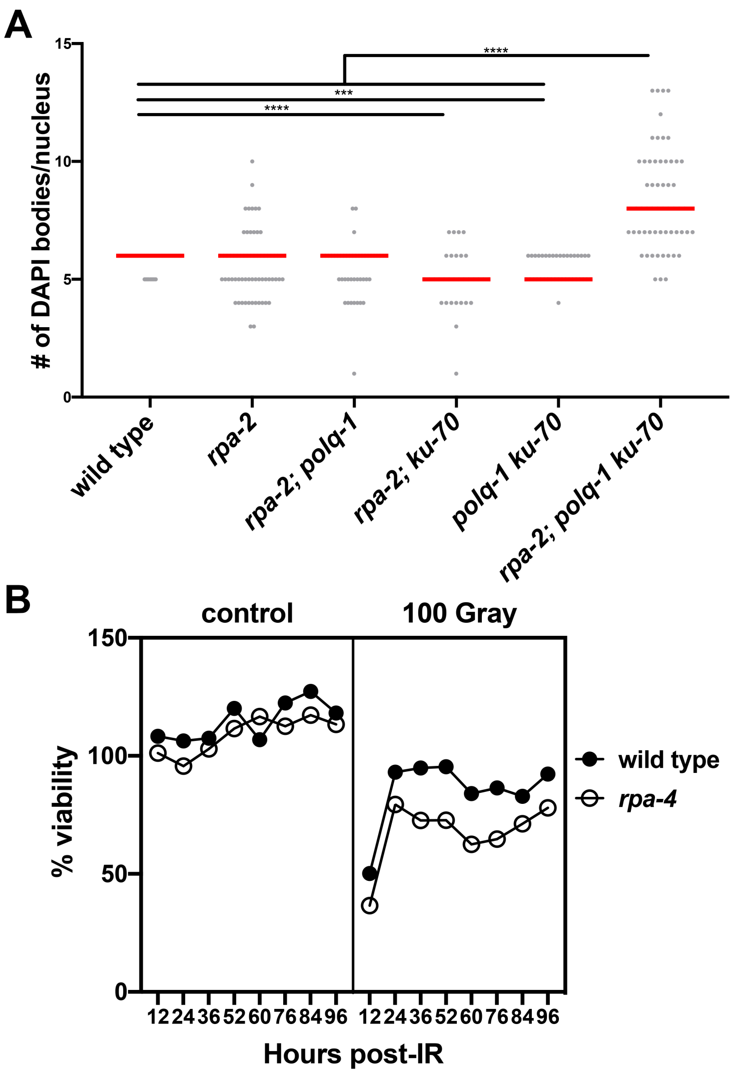


**Figure S4: DAPI body phenotypes of various DSB repair mutants in combination with *rpa-2,* and *rpa-4* sensitivity to gamma-IR with respective controls.**

A) Number of DAPI bodies counted in diakinesis -1 nuclei. B) Percent of laid eggs that hatched at different time points after exposure of adults to gamma-IR.

A

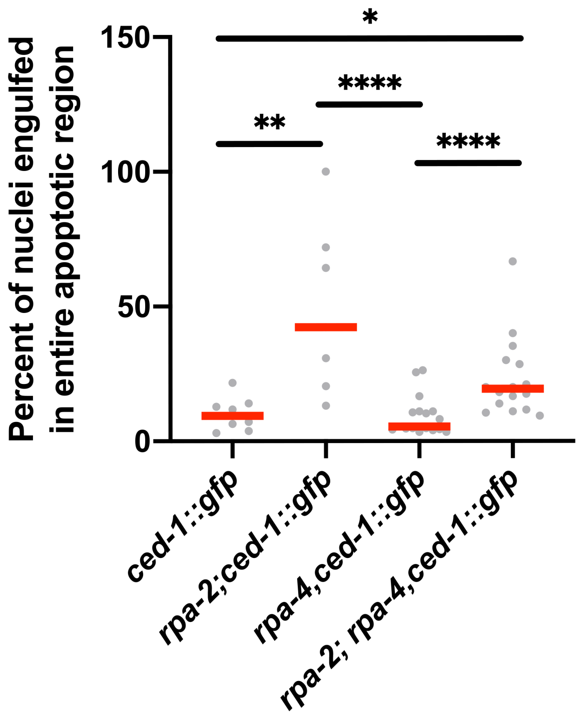


**Figure S5: Comparison of percent CED-1::GFP engulfed nuclei in RPA subunit mutant backgrounds**. Comparison of the percent of engulfed nuclei from the first marked nucleus to the end of pachytene. Each point represents a single gonad. (Mann-Whitney tests performed, where p-values are represented as ****=<0.0001, **=<0.01, and * =<0.05)

Figure S6

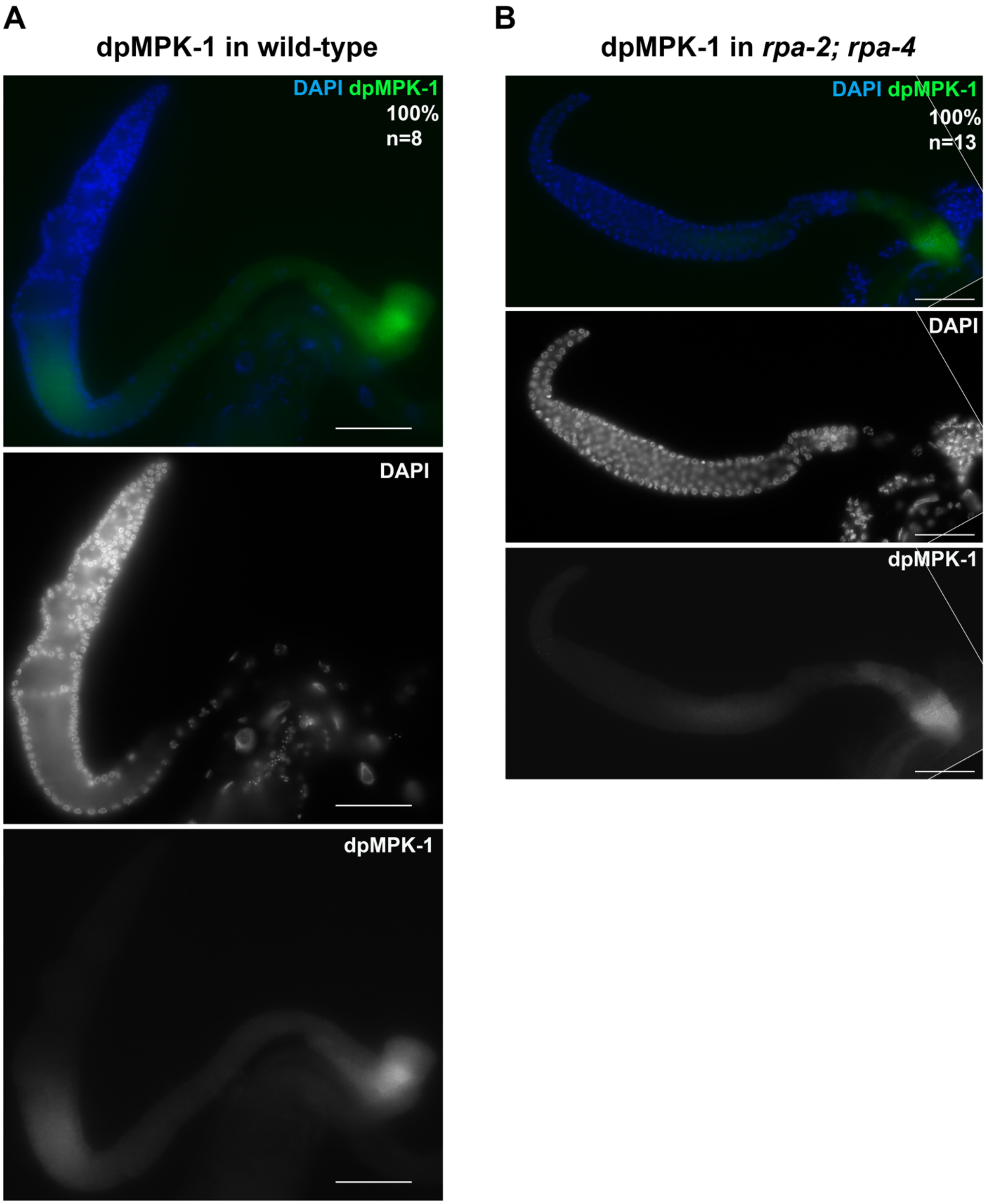


**Figure S6: Representative images of activated MPK-1**
A) Representative images of wild type dpMPK-1 staining. B) Representative images of *rpa-2; rpa-4* double-mutant dpMPK-1 staining. Images taken at 60X magnification. Scale bar= 50 microns.

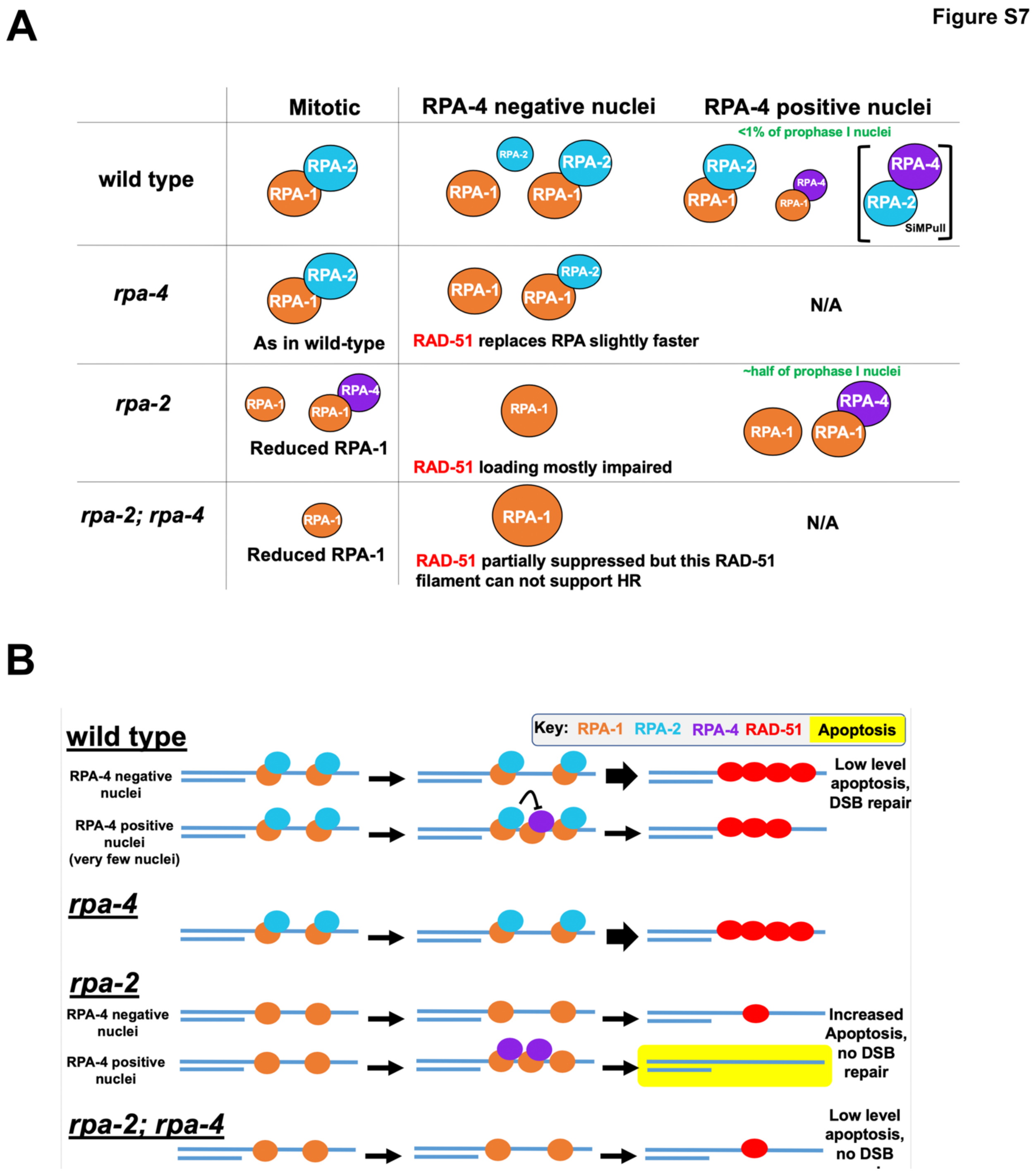


**Figure S7: A model for RPA complexes function**
A) Pictographic description of the cytological and SiMPull data in each genotype, B) Model in context of ssDNA binding and RAD-51 loading
